# Transcript architecture sets the m^6^A landscape: CSTF2 and CSTF2T reshape m^6^A through cleavage-dependent and -independent mechanisms

**DOI:** 10.64898/2026.03.27.714597

**Authors:** Caroline J. Aufgebauer, Theodore M. Nelson, Nadia Houerbi, Seth D. Veenbaas, Matthew Tegowski, Esther Luo, Enakshi Sivasudhan, Michael Goneos, Paul Collier, Jacqueline Proszynski, Krista Ryon, Evan M. Violette, Savanna A. Touré, Kate D. Meyer, Christopher E. Mason, Stacy M. Horner

## Abstract

Alternative RNA processing generates extensive transcript diversity, yet how transcript architecture influences selective m^6^A deposition is incompletely understood. Exon-junction-based models explain where m^6^A is excluded, but a positive determinant of m^6^A accumulation remains undefined. Here, we leverage Zika virus-induced changes in m^6^A deposition to uncover determinants of transcript-selective methylation. By integrating GLORI-seq, native METTL3 RNA immunoprecipitation, and nanopore direct RNA sequencing, we generate a single-nucleotide, isoform-resolved map of m^6^A dynamics during infection. We identify over 2,000 dynamic m^6^A sites, many arising from changes in transcript architecture, and pinpoint proximal polyadenylation sites as positive determinants of m^6^A accumulation. The cleavage stimulation factors CSTF2 and CSTF2T drive this remodeling through two routes: redundant induction of intronic polyadenylation, which converts internal exons into terminal exons that expose DRACH motifs to METTL3, and non-redundant, cleavage-independent recruitment of METTL3 near proximal polyadenylation sites, establishing alternative polyadenylation as a key architectural determinant of the m^6^A landscape.

## Introduction

Post-transcriptional regulation of cellular RNA plays a key role in shaping the host response to viral infection, including infection by *Flaviviridae* viruses, such as Zika virus (ZIKV), dengue virus (DENV), West Nile virus (WNV), and hepatitis C virus (HCV), all of which pose a significant human health burden^1, 2^. Upon infecting host cells, these viruses initiate a complex molecular interplay in which host cells activate antiviral factors to restrict infection, while viruses induce proviral factors that subvert host defenses and promote viral replication^3^. The regulation of these opposing antiviral and proviral responses is critical to both host cell survival and viral propagation.

One mechanism by which host-virus dynamics are regulated is through the RNA modification *N*^6^-methyladenosine (m^6^A)^4^. m^6^A is a prevalent internal mRNA modification that influences nearly all aspects of RNA metabolism. This modification is added to cellular mRNA co-transcriptionally by the m^6^A methyltransferase complex, which consists of the catalytic heterodimer METTL3 and METTL14, as well as additional core proteins, including WTAP^5–7^. METTL3 catalyzes methylation of target RNAs at a specific consensus motif, DRACH (D:G/A/U, R:G/A, H:U/C/A), while METTL14 and WTAP play roles in complex stability and RNA binding^6–13^. Notably, m^6^A is a reversible modification that can be removed by the demethylase enzymes ALKBH5 and FTO^14, 15^. When m^6^A is present on an RNA, specific RNA-binding proteins recognize these m^6^A sites and modulate downstream RNA processing, including splicing, export, translation, and decay^16–20^. Together, the coordinated actions of these m^6^A “writers,” “erasers,” and “readers” establish m^6^A as a regulatory layer that broadly influences RNA metabolism, including during viral infection.

During viral infection, this regulatory layer is engaged to adjust the expression of transcripts encoding proviral or antiviral host factors^21^. We previously found that *Flaviviridae* infection selectively alters m^6^A on a defined set of 51 host transcripts and that these m^6^A changes affected splicing and translation of *CIRBP* and *RIOK3*, two cellular transcripts whose encoded proteins regulate infection^22^. While conserved innate immune signaling and ER stress pathways were sufficient to drive *RIOK3* and *CIRBP* m^6^A changes, respectively, the molecular mechanisms underlying selective m^6^A remodeling of specific host transcripts were not established^22^.These observations raise a broader unresolved question: how specific DRACH motifs are selected for methylation in a given cellular state. Recent work has identified the exon junction complex (EJC) as one such determinant: EJCs limit METTL3 access within ∼150 nucleotides of exon-exon junctions, and this range is sufficient to suppress m^6^A in average-length internal exons but not in long internal or terminal exons^23–25^. The exon–intron boundary itself further represses m^6^A at adjacent exon regions through mechanisms that remain incompletely defined^26^. Together, these mechanisms define where m^6^A is *excluded*, but a positive determinant of where m^6^A accumulates, particularly within long terminal exons, has remained unclear.

Recent transcriptome-wide studies using nanopore direct RNA sequencing (DRS) have revealed that m^6^A is frequently deposited in an isoform-specific manner, with the same genomic adenosine exhibiting different methylation levels across transcript isoforms^27–29^. These observations suggest that additional features of transcript architecture influence m^6^A deposition. One major driver of transcript isoform diversity is alternative polyadenylation (APA)^30^, a key RNA processing mechanism in which transcripts are cleaved and polyadenylated at alternative sites to generate mRNAs with distinct 3′ ends. APA within terminal exons can influence RNA polymerase II (RNAPII) transcription dynamics and remodel 3′UTR architecture where m^6^A preferentially accumulates, potentially affecting the accessibility of nascent transcripts to the m^6^A machinery^31–33^. Intronic polyadenylation (IPA) generates alternative terminal exons by utilizing poly(A) sites in intronic regions, thereby reshaping the spatial relationship between DRACH motifs and splice junctions. Notably, viral infection can induce widespread changes in host APA^34, 35^, raising the possibility that infection-driven remodeling of transcript architecture contributes to selective m^6^A deposition.

Here, we used ZIKV infection to perturb the m^6^A epitranscriptome and uncover positive determinants of m^6^A deposition. By integrating GLORI-seq (glyoxal and nitrite-mediated deamination of unmethylated adenosine sequencing) for site-specific m^6^A quantification^36, 37^, METTL3 RNA immunoprecipitation and sequencing (RIP-seq) to map METTL3 binding^38^, and nanopore direct RNA sequencing (DRS) to resolve m^6^A sites on individual transcript isoforms^39^, we generated a transcript- and site-specific stoichiometric map of m^6^A dynamics, as well as the METTL3 RNA binding profile, in uninfected and ZIKV-infected cells. This comprehensive analysis revealed previously unrecognized m^6^A-altered host transcripts and isoform-specific regulation of m^6^A during infection. Importantly, we found that m^6^A sites are enriched and highly methylated immediately upstream of proximal polyadenylation sites across the transcriptome independently of infection, identifying proximal polyadenylation as a positive determinant of m^6^A accumulation. We further identified the cleavage stimulation factors CSTF2 and CSTF2T as drivers of ZIKV-induced m^6^A remodeling, acting through both cleavage-dependent intronic polyadenylation and cleavage-independent recruitment of METTL3 near proximal polyadenylation sites. Together, these findings establish transcript architecture shaped by alternative polyadenylation as a key determinant of where m^6^A accumulates, with CSTF2 and CSTF2T as one route by which it is reshaped during infection.

### GLORI-seq reveals selective remodeling of host m^6^A sites by ZIKV

In previous work, we used methylated RNA immunoprecipitation and sequencing (meRIP-seq) to identify 54 common dynamic m^6^A peaks in 51 transcripts following infection by the *Flaviviridae* viruses ZIKV, DENV, WNV, and HCV^22^. However, the exact modified adenosines and the absolute stoichiometries of m^6^A at these sites remained unknown due to the limited resolution and quantitative capacity of meRIP^40^. GLORI-seq alleviates these challenges by leveraging chemical deamination of unmodified adenosines while sparing m^6^A-modified adenosines, enabling their detection at single-nucleotide resolution by RNA sequencing. As such, this method precisely identifies individual m^6^A sites and enables relative quantification of modification stoichiometry across transcripts^36^. Thus, we performed GLORI-seq on poly(A)-enriched RNA from mock- and ZIKV-infected Huh7 cells, a model cell line that supports robust ZIKV infection^41^, at 48 hours post-infection (**Figure 1A**). We identified 67,130 and 66,880 m^6^A sites in mock and ZIKV-infected cells (m^6^A level≥10%, p_adj_<0.05), respectively, with a median m^6^A level of ∼26% in both conditions and reproducible m^6^A levels between three biologically independent replicates (**Figure S1A-C; Table S1**). Principal component analysis revealed clustering by infection status and of biological replicates, indicating high reproducibility and minimal technical variability (**Figure S1D**). Similar to previous studies^10, 11, 36^, m^6^A sites were distributed near stop codons and within 3′ untranslated regions (UTRs) in both conditions, and the consensus m^6^A motif DRACH (D = A, G, or U; R = A or G; H = A, C, or U) was enriched at these sites, confirming the work of others that GLORI-seq captures known features of m^6^A (**Figure S1E-F**)^36^.

**Figure 1.**
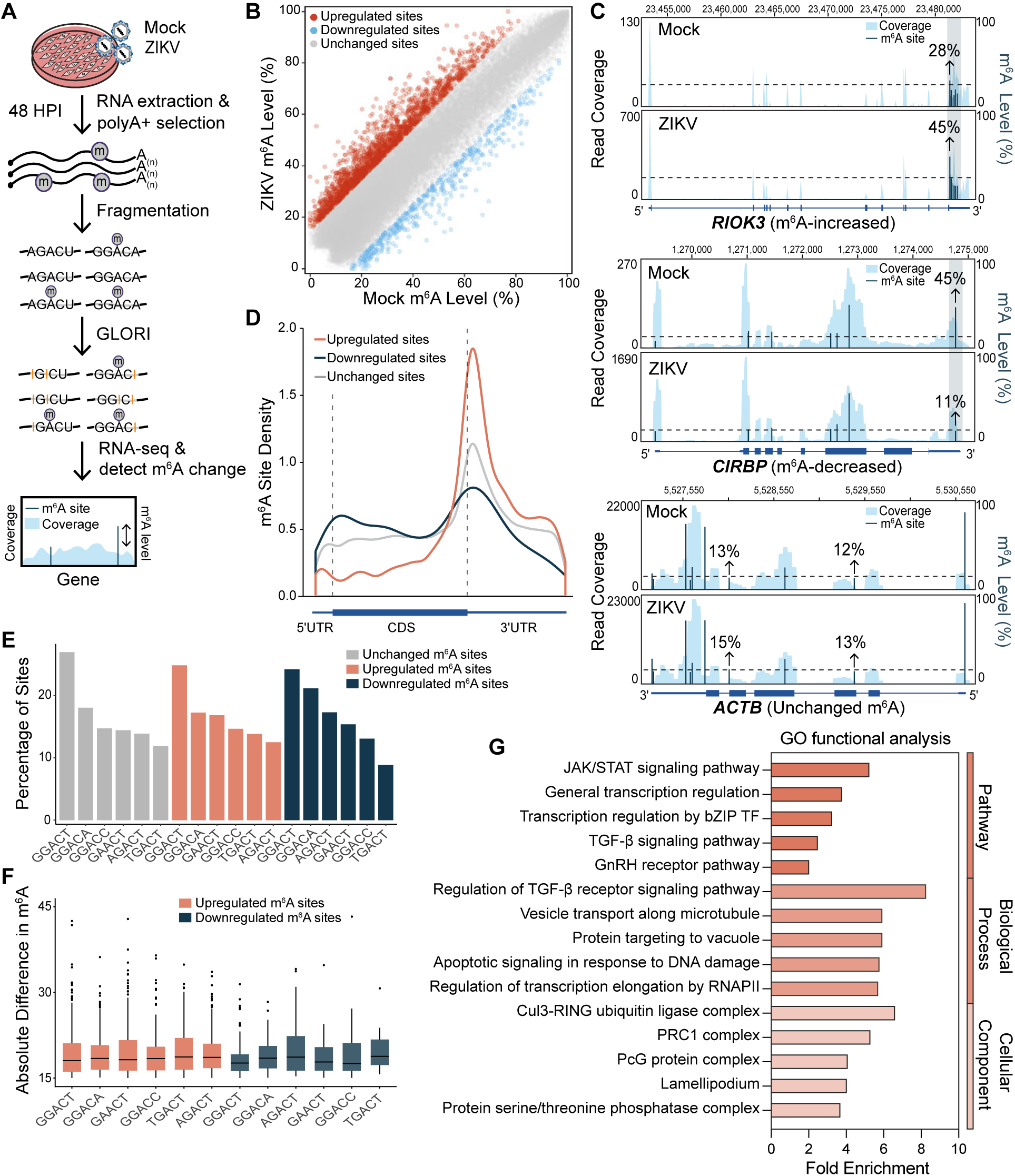
ZIKV infection alters m^6^A levels on cellular transcripts. **A)** Experimental workflow of GLORI-seq protocol to identify differential m^6^A following ZIKV infection (MOI 1, 48 hpi, n=3). **B)** Scatter plot showing ZIKV-induced changes in m^6^A levels of individual sites. Orange and blue dots represent upregulated (n=2004) and downregulated (n=502) m^6^A sites during ZIKV infection, respectively. **C)** Gene track plots of upregulated m^6^A in *RIOK3* (upper panel), downregulated m^6^A in *CIRBP* (middle panel), and unchanged m^6^A in *ACTB* (lower panel). **D)** Metagene profiles showing the distributions of upregulated, downregulated, and unchanged m^6^A sites across the mRNA molecule. **E)** The top 6 motifs across upregulated, downregulated, and unchanged m^6^A sites and the proportion of m^6^A sites they contain. **F)** The absolute difference in m^6^A level of upregulated and downregulated m^6^A sites across the top 6 motifs. **G)** PANTHER GO enrichment analysis for genes with upregulated m^6^A sites using a binomial test with FDR correction (FDR corrected p < 0.05).

We next investigated the dynamic regulation of m^6^A sites upon ZIKV infection. Using a ± 15% absolute methylation level change threshold (|Δm^6^A ratio|≥0.15, p<0.05)^36^, we identified 2,004 upregulated and 502 downregulated m^6^A sites upon ZIKV infection, corresponding to 1,413 and 450 genes, respectively (**Figure 1B**). Thus, approximately 2% of m^6^A sites are dynamic in response to ZIKV infection, revealing a highly selective modulation of m^6^A levels. To benchmark GLORI-seq against our previously published meRIP-seq data (**Note S1**), we examined two functionally validated m^6^A peaks: one upregulated on *RIOK3* and one downregulated on *CIRBP*^22^. GLORI-seq recapitulated these dynamic changes, while m^6^A sites in the unchanged control *ACTB* remained stable upon infection (**Figure 1C**). In summary, GLORI-seq uncovered thousands of dynamic m^6^A sites upon ZIKV infection that were previously undetectable by meRIP-seq.

We next examined the characteristics of dynamic m^6^A sites to identify distinguishing features associated with selective m^6^A regulation. Upregulated m^6^A sites displayed absolute m^6^A level increases ranging from 15% to 53%, with a median change of 18% (**Table S2**). Downregulated m^6^A sites showed comparable changes ranging from −15% to −46%, with a median change of −18% (**Table S2**). These upregulated and downregulated m^6^A sites differed slightly in their distribution patterns across the mRNA molecule. Specifically, we observed enrichment of upregulated m^6^A sites around the stop codon, similar to unchanged m^6^A sites, while downregulated m^6^A sites had enrichment around the stop codon as well as slight enrichment at the 5′UTR (**Figure 1D**). The top preferred DRACH motifs, however, remained highly consistent across conditions, with only slight differences in their ranking (**Figure 1E**). Further, the absolute difference in m^6^A level of upregulated and downregulated m^6^A sites did not show any striking differences across motifs, suggesting that motif usage is not a major driver of dynamic m^6^A regulation (**Figure 1F**). Taken together, these data indicate that the canonical features of m^6^A modification are not broadly altered during ZIKV infection.

m^6^A has been previously shown to regulate transcripts involved in infection and innate immune signaling^4, 42–44^. To further explore if m^6^A-altered genes during ZIKV infection shared these features, we performed functional annotation of m^6^A-increased and m^6^A-decreased genes across PANTHER Gene Ontology (GO) categories: Pathway, Biological Process, and Cellular Component (**Figure 1G**). m^6^A-decreased genes showed no statistically significant pathway enrichment (FDR>0.05); thus, our subsequent analyses in this manuscript focus on m^6^A-increased genes. These genes are enriched for JAK/STAT and TGF-β signaling pathways, key immune regulatory pathways implicated in viral infection^45–48^. Consistent with this, regulation of the TGF-β signaling pathway emerged as the top enriched biological process, along with other processes typically manipulated during viral infection, such as vesicle transport along microtubules, apoptotic signaling in response to DNA damage, and regulation of RNAPII transcription elongation (**Figure 1G**). Given that ZIKV targets specific host cell machinery to subvert host cell defenses and facilitate its replication^47, 49–51^, we next examined which cellular components were enriched among m^6^A-increased genes. The top enriched cellular component is the Cul3-RING ubiquitin ligase complex, which ZIKV hijacks to degrade STAT2, a key signaling intermediate of the JAK/STAT signaling pathway^48^. Together, these findings reveal that ZIKV infection alters m^6^A levels on genes encoding key regulators of host antiviral responses.

### m^6^A-altered genes are differentially targeted by METTL3 in an isoform-specific manner during ZIKV infection

As the expression levels of the m^6^A machinery are unchanged upon *Flaviviridae* infection^22^, we hypothesized that altered m^6^A levels during ZIKV infection are driven by differential RNA targeting of METTL3. To test this, we performed native RNA immunoprecipitation with an anti-METTL3 antibody to enrich METTL3-bound RNA in mock- and ZIKV-infected Huh7 cells at 48 hours post-infection (**Figure 2A**). Given that m^6^A can be regulated in an isoform-specific manner^39, 52^, we performed immunoprecipitation on full-length RNA and implemented a transcript-level analysis of METTL3 RNA binding, enabling detection of isoform-level regulatory changes that are typically obscured by fragmentation and aggregated gene-level approaches. Variance-stabilized transcript-level counts were clustered by assay and infection status, with replicates tightly grouped, indicating high reproducibility and minimal technical variability (**Figure S2A**). Before evaluating differential binding of METTL3 upon infection, we first identified transcripts that are truly METTL3-bound in mock and infected conditions independently. In total, we identified 9,521 and 10,551 transcripts (5,023 and 6,392 genes) significantly enriched in the METTL3 RIP samples relative to the input samples in mock- and ZIKV-infected cells, respectively (**Figure S2B, Table S3**). We next asked to what extent METTL3 binding corresponded to m^6^A modification by comparing transcripts identified in METTL3 RIP-seq and GLORI-seq analyses. For these cross-assay comparisons, we restricted analysis to genes that passed coverage thresholds in both datasets, ensuring that lack of overlap reflected true biological differences rather than insufficient read coverage. Consistent with the established role of METTL3 in installing m^6^A on mRNA^5, 6^, 91% and 92% of METTL3-bound transcripts were also identified as m^6^A-modified by GLORI-seq in mock- and ZIKV-infected conditions, respectively (**Figure S2C**).

**Figure 2.**
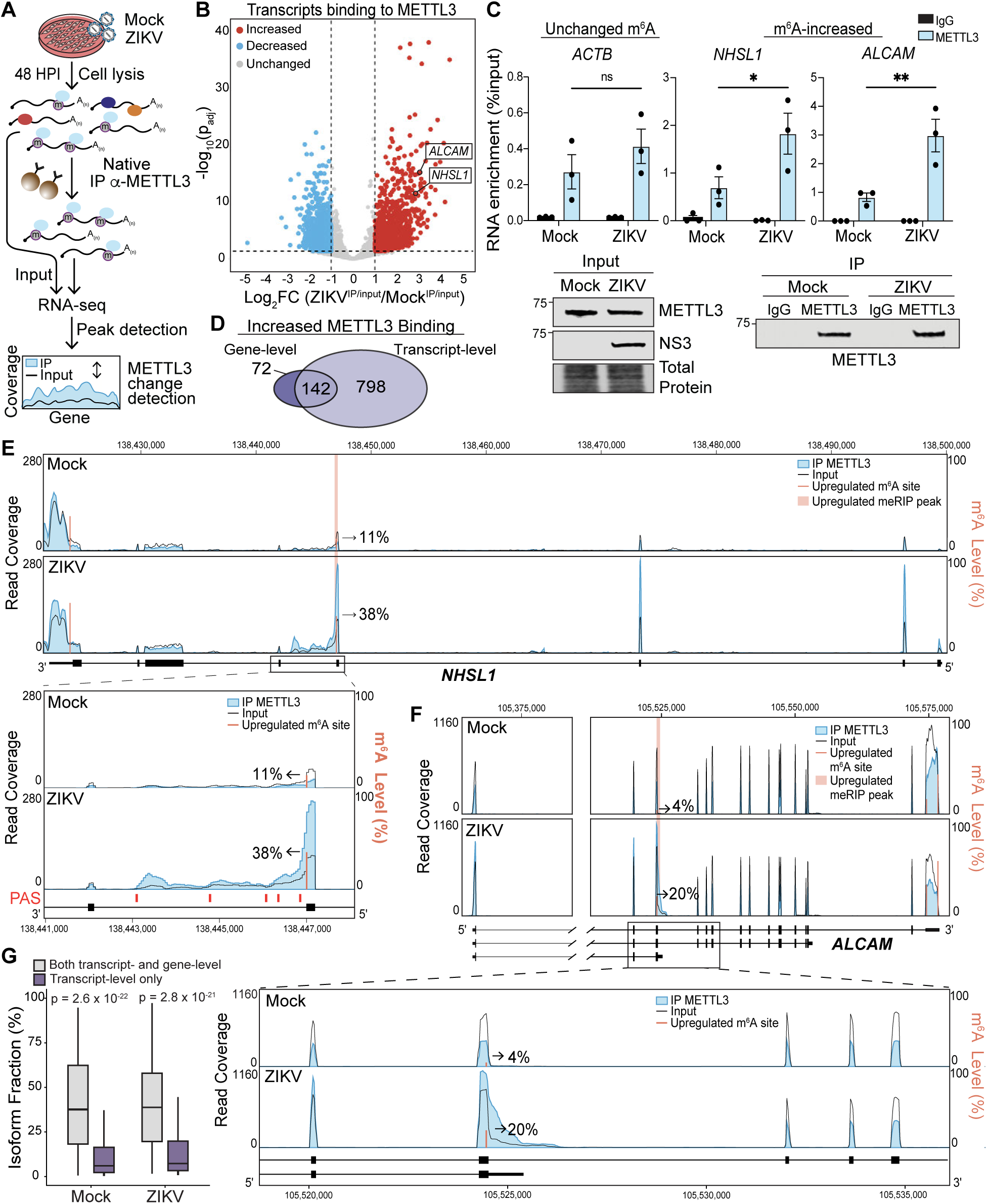
ZIKV infection induces differential METTL3 targeting to cellular transcripts. **A)** Experimental workflow of METTL3 RIP-seq protocol to identify differential METTL3 binding following ZIKV infection (MOI 1, 48 hpi, n=4). **B)** Volcano plot showing ZIKV-induced changes in METTL3 binding to transcripts. Red and blue dots represent increased (Log_2_FC ≥ 1 and p_adj_ < 0.05) and decreased (Log_2_FC ≤ −1 and p_adj_ < 0.05) METTL3 binding during ZIKV infection, respectively. **C)** (Top) METTL3 RIP RT-qPCR analysis of relative METTL3 binding of transcripts with ZIKV-altered METTL3 binding or control (*ACTB*) in ZIKV-infected (MOI 1, 48 hpi) Huh7 cells. n = 3 biologically independent experiments, with bars indicating mean values and error bars showing standard errors of the mean. *p < 0.05, **p < 0.01, or not significant (ns), determined by two-way ANOVA followed by Šidák’s multiple comparison test. (Bottom) Representative immunoblots of inputs and immunoprecipitated lysates from n=3 biologically independent experiments. **D)** Venn diagram showing the overlap of genes detected to have increased METTL3 binding in the gene-level analysis (STAR-based workflow) and the transcript-level analysis (Salmon-based workflow). **E)** Gene track plot of METTL3 binding in *NHSL1*, along with quantification of m^6^A levels at specific sites, as determined by GLORI-seq, and annotation of PAS sites from the PolyASite database^53–55^. **F)** Gene track plot of METTL3 binding in *ALCAM*, along with quantification of m^6^A levels at specific sites, as determined by GLORI-seq. **G)** Boxplot comparing the isoform fractions (percentage of a gene’s total expression contributed by each isoform) for isoforms from genes with increased METTL3 binding at both the transcript- and gene-level analysis and exclusively at the transcript-level.

We next examined how ZIKV infection alters METTL3 binding to cellular transcripts. We found that infection led to increased METTL3 binding on 1,065 transcripts (940 genes) and decreased METTL3 binding on 1,506 transcripts (1,127 genes) (**Figure 2B, Table S4)**. Among the top genes with increased METTL3 binding and upregulated m^6^A sites were *ALCAM* and *NHSL1*, which we also previously found to have upregulated meRIP peaks^22^, and these peaks directly overlap upregulated m^6^A sites that we detected by GLORI-seq **(Table S2, Table S4, Figure S2D)**. By using METTL3 RIP RT-qPCR, we validated increased METTL3 binding on *ALCAM* and *NHSL1*, while the unchanged m^6^A-modified control *ACTB* had no change in METTL3 binding (**Figure 2C**). Of note, 85% of genes with increased METTL3 binding in the transcript-level analysis were overlooked by gene-level analysis, including *ALCAM* and *NHSL1* (**Figure 2D, Table S54-5**). Similarly, 74% of genes with decreased METTL3 binding at the transcript-level analysis were missed at the gene-level (**Figure S2E, Table S54-5**). Because gene-level analyses aggregate reads across all transcripts of a gene, these results suggest that many differential METTL3 binding events occur on specific transcript isoforms rather than uniformly across all transcript isoforms of a gene. Indeed, we observed isoform-specific METTL3 binding patterns in the gene tracks for *ALCAM* and *NHSL1* (**Figure 2E-F**). Specifically, *NHSL1* appears to have an unannotated shorter isoform, in which the fourth exon becomes the terminal exon and is extended with a 3′UTR (**Figure 2E**). This isoform is consistent with usage of an annotated alternative poly(A) site (PAS) from the PolyASite database^53–55^ that is located downstream of the fourth exon (**Figure 2E**). ZIKV infection induces strong enrichment of METTL3 over the genomic region corresponding to the short *NHSL1* isoform, and METTL3 binding drops sharply beyond this isoform (**Figure 2E**). *ALCAM* also encodes multiple alternative isoforms, the shortest of which has increased read coverage upon ZIKV infection (**Figure 2F**). Notably, we found that upon ZIKV infection, METTL3 is only enriched over the genomic region corresponding to the short isoform and lacks detectable binding beyond this isoform, consistent with isoform-specific binding (**Figure 2F**). Together, these data reveal that ZIKV infection differentially targets METTL3 to a subset of m^6^A-altered genes in an isoform-specific manner.

To investigate why many METTL3-altered genes detected by the transcript-level analysis were overlooked in the gene-level analysis, we next examined the relative abundance of transcript isoforms. Using IsoformSwitchAnalyzeR^56^, we calculated isoform fractions, defined as the percentage of a gene’s total expression contributed by each isoform. We found that isoforms from genes in which increased METTL3 binding was exclusively detected in the transcript-level analysis exhibited significantly lower isoform fractions than isoforms from genes detected in both gene- and transcript-level analyses (**Figure 2G**). This suggests that many METTL3-altered transcript isoforms are relatively low-abundance isoforms. Because gene-level quantification aggregates sequencing reads across all transcript isoforms of a gene, binding changes restricted to these minor isoforms can be diluted by more highly expressed isoforms and therefore remain undetected in gene-level analyses. Overall, these findings demonstrate that a substantial portion of METTL3 binding changes during ZIKV infection occur at the transcript isoform level, and that minor isoform populations can serve as previously unrecognized targets of differential m^6^A regulation.

### m^6^A is regulated in an isoform-specific manner during ZIKV infection

Given that METTL3 binds specific RNA isoforms, we next asked whether m^6^A is regulated in an isoform-specific manner during ZIKV infection. Nanopore DRS enables full-length, single-molecule sequencing of native transcripts, allowing m^6^A modifications to be directly assigned to individual transcript isoforms^27, 29^. We therefore performed DRS on poly(A)-enriched RNA from mock- and ZIKV-infected Huh7 cells at 48 hours post-infection (**Figure 3A**). We obtained an average of approximately 17.3 million primary aligned reads per sample (range: 13.1-21.1 million), with a mean read length of 1.36 kb, providing sufficient depth for m^6^A stoichiometry estimates at the transcript level. Reads were base-called and then m^6^A sites were identified using Dorado^57^. Using principal component analysis, we found that our samples showed clear separation by infection status and clustering of biological replicates, indicating high reproducibility and minimal technical variability (**Figure S3A**). Applying an FDR cutoff of 5%, a minimum coverage of 15 reads per site, and a minimum modification rate of 10%, we identified 85,048 and 95,493 m^6^A sites in mock- and ZIKV-infected cells, respectively, with strong agreement across our three biologically independent replicates (**Figure S3B-C**). DRS-identified m^6^A sites exhibited canonical m^6^A features, including preferential localization near stop codons and within 3′UTRs, as well as enrichment of the DRACH consensus motif (**Figure S3D-E**). We next compared m^6^A sites predicted by DRS with those detected using GLORI-seq (**Figure 3B-C**). To avoid confounding biological discordance with differences in sequencing depth, we restricted comparison to genes that passed coverage thresholds in both datasets, ensuring that lack of overlap reflected true discrepancies in m^6^A detection rather than insufficient read coverage. We found 62-74% overlap between GLORI-seq and DRS m^6^A sites, as well as strong agreement (R^2^=92-93%) in m^6^A stoichiometry at shared sites (**Figure 3B-C**). Together, these results demonstrate robust and reproducible m^6^A detection across orthogonal sequencing platforms.

**Figure 3.**
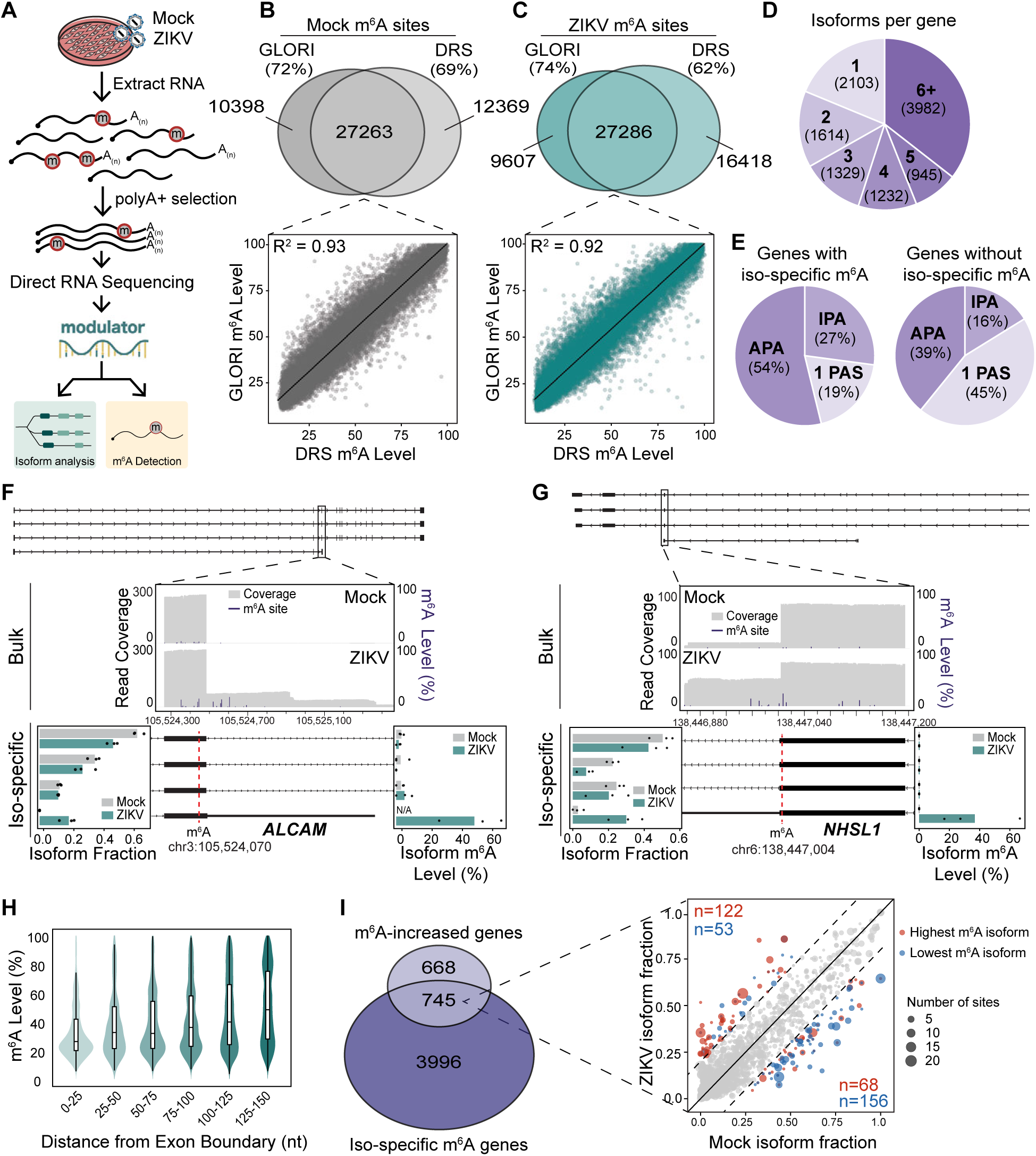
m^6^A is regulated in an isoform-specific manner during infection. **A)** Experimental workflow of DRS protocol to identify m^6^A at isoform-level resolution following ZIKV infection (MOI 1, 48 hpi, n=3). **B)** (Top) Venn diagram showing the overlap of sites detected to be m^6^A-modified in GLORI-seq and DRS in the mock condition. (Bottom) Scatter plot showing the predicted m^6^A levels of shared sites in GLORI and DRS. **C)** (Top) Venn diagram showing the overlap of sites detected to be m^6^A-modified in GLORI-seq and DRS in ZIKV condition. (Bottom) Scatter plot showing the predicted m^6^A levels of shared sites in GLORI and DRS. **D)** Pie chart of number of isoforms per gene detected by Modulator. **E)** (Left) Pie chart of percentage of APA classes of genes with isoform-specific m^6^A and METTL3 binding. (Right) Pie chart of percentage of APA classes of genes without isoform-specific m^6^A. **F)** Representative gene track plot of *ALCAM*, along with quantification of m^6^A levels at specific sites and quantification of isoform fractions, as determined by DRS and Modulator. **G)** Representative gene track plot of *NHSL1*, along with quantification of m^6^A levels at specific sites and quantification of isoform fractions, as determined by DRS and Modulator. **H)** Comparison of transcriptome-wide m^6^A levels of sites at increasing distances from an exon-exon junction. **I)** (Left) Venn diagram showing the overlap of m^6^A-increased genes in GLORI-seq and genes with isoform-specific m^6^A in DRS. (Right) Scatter plot comparing isoform fractions in mock and ZIKV for isoforms from genes with ZIKV-induced increased m^6^A and isoform-specific m^6^A. For each site, the isoform with the highest m^6^A level in ZIKV (“Highest m^6^A isoform”, red) and the isoform with the lowest m^6^A level (“Lowest m^6^A isoform”, blue) were identified. Each point represents a unique isoform collapsed across all associated sites, with point size indicating the number of contributing m^6^A sites. Dashed lines indicate ± 0.2 isoform fraction change.

To define a transcript-specific map of m^6^A, we applied Modulator^58^, a novel algorithm that probabilistically assigns truncated DRS reads to transcript fragments to identify transcript-related epitranscriptomic site-level stoichiometric variation. Modulator enables rigorous disambiguation of isoform-specific m^6^A in infection contexts where APA generates novel isoforms absent from reference annotations. We detected multiple isoform fragments for the majority of genes, with a median of four unique isoform constructions per gene (**Figure 3D**). We then compared methylation levels of m^6^A sites across these different RNA isoform fragments from the same gene, revealing 13,852 m^6^A sites that exhibit isoform-specific m^6^A (p<0.05, absolute m^6^A level difference≥10%) (**Table S6**). Notably, genes with isoform-specific increases in METTL3 binding strongly overlapped those with isoform-specific m^6^A, including *ALCAM* and *NHSL1* (**Figure S3F**). These preferentially modified and METTL3-bound isoforms were enriched for products of intronic polyadenylation (IPA) (**Figure 3E**). For both *ALCAM* and *NHSL1*, the same genomic adenosine was m^6^A-modified when located in a terminal exon but unmodified when present within an internal exon of an alternative isoform (**Figure 3F-G**), consistent with prior work showing that m^6^A deposition is suppressed near exon-exon junctions^23–26^. Accordingly, transcriptome-wide analysis revealed m^6^A depletion within 0-100 nucleotides of exon-exon junctions (**Figure 3H**), supporting a model in which IPA generates new terminal exons that escape this junction-proximal suppression and thereby acquire m^6^A. Additionally, the highly modified IPA isoforms of *ALCAM* and *NHSL1* were upregulated upon ZIKV infection, accounting for the observed increase in bulk m^6^A of these genes (**Figure 3F-G**). Together, these findings suggest that a subset of ZIKV-induced m^6^A changes arise from changes in exon architecture, whereby IPA generates terminal exons that are permissive for m^6^A modification.

Because GLORI-seq detects changes in bulk m^6^A level without resolving transcript isoforms, we next asked how many GLORI-detected m^6^A-increased genes exhibit isoform-specific methylation patterns. We found that 53% of these genes with increased m^6^A contain isoform-specific m^6^A in our DRS data (**Figure 3I, left**). Within this overlapping gene set, isoforms that increased in abundance during ZIKV infection preferentially carried the highest m^6^A levels at a given site, whereas isoforms that decreased in abundance carried the lowest m^6^A levels (**Figure 3I, right**). Together, these results support a model in which some apparent increases in bulk m^6^A levels arise not from uniform remodeling but from ZIKV-induced shifts in isoform usage toward highly methylated isoforms and away from weakly methylated ones. The IPA-driven mechanism described above represents one example of this broader pattern.

### The cleavage stimulation factors CSTF2 and CSTF2T promote intronic polyadenylation of *ALCAM* and *NHSL1* and unmask DRACH motifs for m^6^A modification

We next investigated the mechanism driving upregulation of the highly methylated short isoforms of *ALCAM* and *NHSL1* upon ZIKV infection. Differential isoform expression can be mediated by alternative splicing or APA^30, 59, 60^. To identify factors driving these isoform switches, we analyzed ENCODE enhanced crosslinking and immunoprecipitation (eCLIP) data^61^ from HepG2 cells, another hepatoma-derived liver cell line, for RNA-binding proteins enriched at the relevant 3′ regions. This revealed that the cleavage stimulation factors CSTF2 and CSTF2T bind downstream of the terminal exons of these isoforms of *ALCAM* and *NHSL1* (**Figure S4A-B**), suggesting a role in altered 3′ end processing. CSTF2 and its paralog CSTF2T (CSTF2/T) act redundantly to drive cleavage at noncanonical poly(A) sites, including proximal poly(A) sites that shorten 3′UTRs or intronic poly(A) sites that generate truncated transcripts^62–64^. Thus, we hypothesized that upregulation of the short isoforms of *ALCAM* and *NHSL1* during infection is driven by CSTF2/T-mediated cleavage at intronic poly(A) sites. Using RT-qPCR primers specific to the short and long isoforms of *ALCAM* and *NHSL1*, we confirmed that ZIKV infection increases the abundance of the short-isoforms, while decreasing the abundance of the long-isoforms of both *ALCAM* and *NHSL1,* consistent with the isoform shift observed by DRS (**Figure 4A-B**).

**Figure 4.**
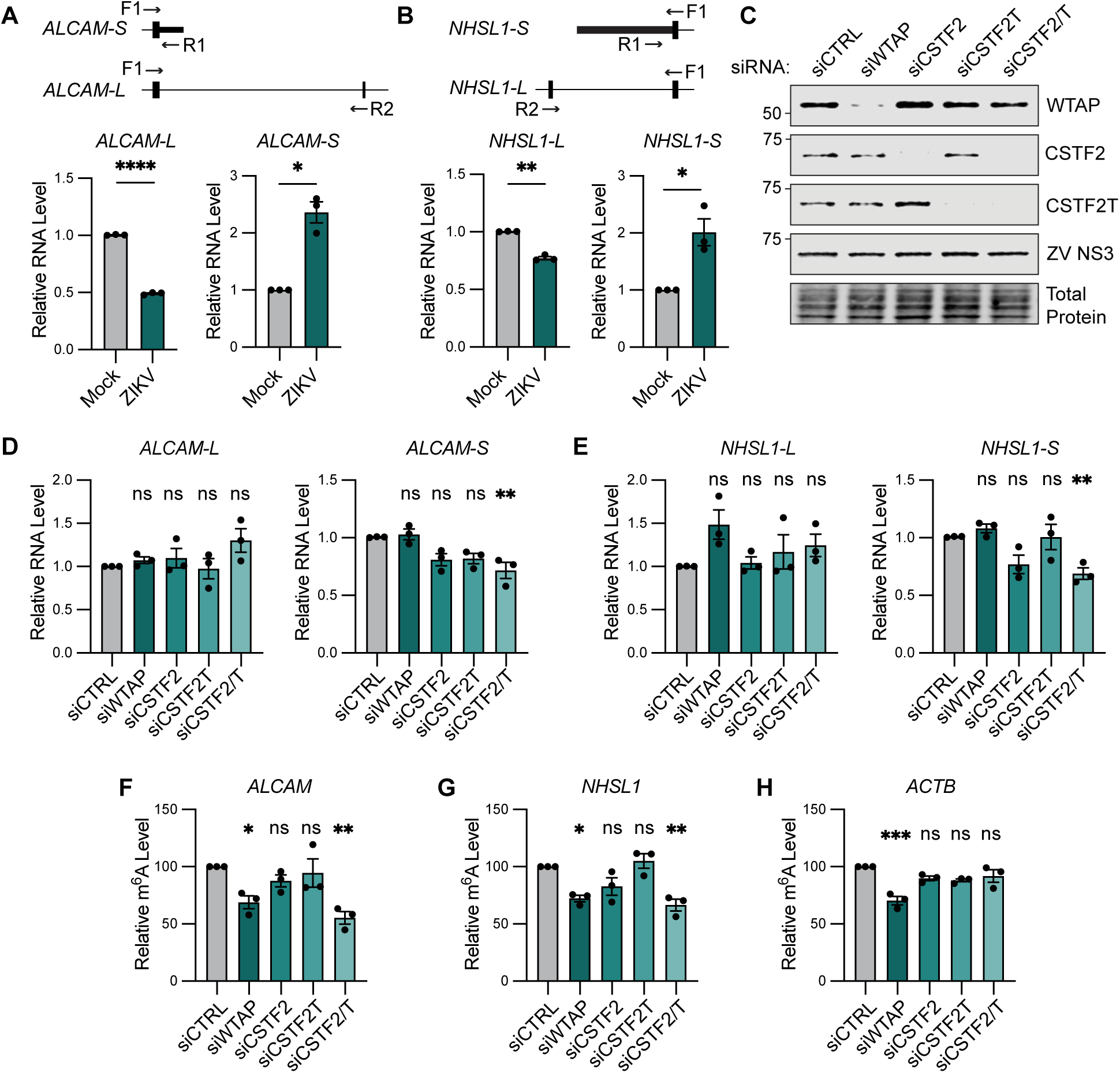
CSTF2 and CSTF2T regulate intronic polyadenylation and m^6^A modification of *ALCAM* and *NHSL1* during ZIKV infection. **A)** and **B)** (Top) Schematic of primers designed to specifically amplify short and long isoforms of *ALCAM* (A) and *NHSL1* (B). (Bottom) RT-qPCR analysis (relative to RPL30) of RNA expression of *ALCAM* and *NHSL1* isoforms in Huh7 cells infected with ZIKV (48 hpi, MOI 0.5). **C)** Immunoblot analysis of lysates from Huh7 cells treated with the indicated siRNA followed by ZIKV infection (48 hpi, MOI 0.5). Representative immunoblot from n=3 biologically independent experiments. **D)** and **E)** RT-qPCR analysis (relative to *RPL30*) of RNA expression of *ALCAM* (D) and *NHSL1* (E) isoforms in Huh7 cells treated with the indicated siRNA followed by ZIKV infection (48 hpi, MOI 0.5). **F)** and **G)** meRIP-RT-qPCR analysis of relative m^6^A level of *ALCAM* (F) and *NHSL1* (G) in Huh7 cells treated with the indicated siRNA followed by ZIKV infection (48 hpi, MOI 0.5). **H)** meRIP-RT-qPCR analysis of relative m^6^A level of *ACTB* in Huh7 cells treated with the indicated siRNA followed by ZIKV infection (48 hpi, MOI 0.5). For all panels, n = 3 biologically independent experiments, with bars indicating mean and error bars showing the standard error. *p < 0.05, **p < 0.01, ***p < 0.001, or not significant (ns) as determined by Welch’s t-test (**A** and **B**) and one-way analysis of variance (ANOVA) with Dunnett’s multiple comparison test (**D**-**H**).

Given that CSTF2 and CSTF2T are predicted to bind downstream of the short-isoform terminal exons of *ALCAM* and *NHSL1*, we next asked whether CSTF2/T are functionally required for the ZIKV-induced isoform switch by performing individual and combined siRNA knockdown followed by RT-qPCR. As some studies have shown examples of APA events regulated by m^6^A itself^65, 66^, we modulated m^6^A levels by siRNA-mediated depletion of WTAP, an essential regulatory subunit of the m^6^A methyltransferase complex^7^. Knockdown efficiency was confirmed by immunoblot, and ZIKV NS3 protein levels were unchanged across all siRNA conditions (**Figure 4C**), indicating that depletion of CSTF2 and CSTF2T did not impair viral replication. Consistent with this, depletion of CSTF2 and CSTF2T did not alter ZIKV infectious titer (**Figure S4C**). WTAP depletion did not affect expression of the short or long isoforms of *ALCAM* and *NHSL1* (**Figure 4D-E**). In contrast, combined depletion of CSTF2 and CSTF2T significantly reduced expression specifically of the short isoforms, whereas individual depletion of CSTF2 or CSTF2T had no effect, consistent with their known redundancy in APA (**Figure 4D-E**).

To test whether CSTF2/T-driven IPA regulates m^6^A in *ALCAM* and *NHSL1,* we performed meRIP-RT-qPCR following CSTF2 and/or CSTF2T depletion. Combined depletion of both CSTF2 and CSTF2T resulted in significant loss of m^6^A on *ALCAM* and *NHSL1,* at levels comparable to those seen with WTAP depletion, while individual knockdown of either CSTF2 or CSTF2T did not affect m^6^A levels (**Figure 4F-G**). As expected, depletion of CSTF2 and CSTF2T had no effect on m^6^A levels of *ACTB*, which is neither alternatively polyadenylated nor differentially methylated during ZIKV infection (**Figure 4H**). Taken together, these data indicate that changes in bulk m^6^A levels on *ALCAM* and *NHSL1* during ZIKV infection are driven by CSTF2/T-dependent IPA, which converts an internal exon to a terminal exon and thereby renders these short isoforms accessible to METTL3.

We previously showed by meRIP-seq that *ALCAM* and *NHSL1* acquire m^6^A in response to multiple *Flaviviridae* viruses, including DENV, WNV, and HCV^22^. We therefore asked whether these viruses also induce expression of the short isoforms of *ALCAM* and *NHSL1*. Indeed, both DENV and WNV infection upregulated the short isoforms of *ALCAM* and *NHSL1*, similar to ZIKV **(Figure S5A-B**), and HCV showed an upward trend in short isoform expression that did not reach statistical significance (**Figure S5C**). In contrast, transfection of a RIG-I pathogen-associated molecular pattern (PAMP)^67^, which activates RIG-I-mediated innate immune signaling without altering m^6^A on these transcripts^22^, did not induce the short isoforms **(Figure S5D5D-E**). Together, these data indicate that the isoform switch toward short, m^6^A permissive isoforms is a shared feature across these *Flaviviridae* viruses that cannot be explained by innate immune signaling alone.

### CSTF2 and CSTF2T regulate tandem 3′UTR APA and m^6^A on *NR1D2* and *MXD1* through cleavage-independent mechanisms

The data above identify one mechanism by which APA reshapes the m^6^A landscape: IPA generates new terminal exons that escape exon junction-proximal suppression. However, many ZIKV-induced changes in METTL3 binding occur on transcripts that do not undergo IPA (**Figure 3E**), indicating that additional mechanisms for altered m^6^A exist. We therefore examined a second, mechanistically distinct class of APA: tandem 3′UTR shortening, in which proximal poly(A) site usage produces a shorter 3′UTR without altering exon-intron architecture. Unlike the case of IPA, m^6^A changes on transcripts generated by tandem APA cannot be explained by EJC loss alone, raising the question of how m^6^A levels on these transcripts are modulated. We focused on two ZIKV-altered transcripts that exhibit tandem 3′UTR APA, *NR1D2* and *MXD1*. These genes are among the top genes with increased METTL3 binding and upregulated m^6^A sites (**Table S2**; **Table S5).** Notably, *MXD1* was detected to have an upregulated m^6^A peak during ZIKV infection by meRIP-seq^22^, which directly overlaps an upregulated m^6^A site that we detected by GLORI-seq. *MXD1* encodes the MAX dimerization protein 1, a transcriptional repressor that antagonizes MYC-dependent gene expression and has been previously shown to regulate influenza virus-induced cytokine production^68^. *NR1D2* encodes the nuclear receptor REV-ERBΔ, a circadian regulator that modulates innate immune responses^69^. Both *NR1D2* and *MXD1* undergo APA during ZIKV infection, with METTL3 binding preferentially enriched over the shortened 3′UTR region (**Figure 5A-B**).

**Figure 5.**
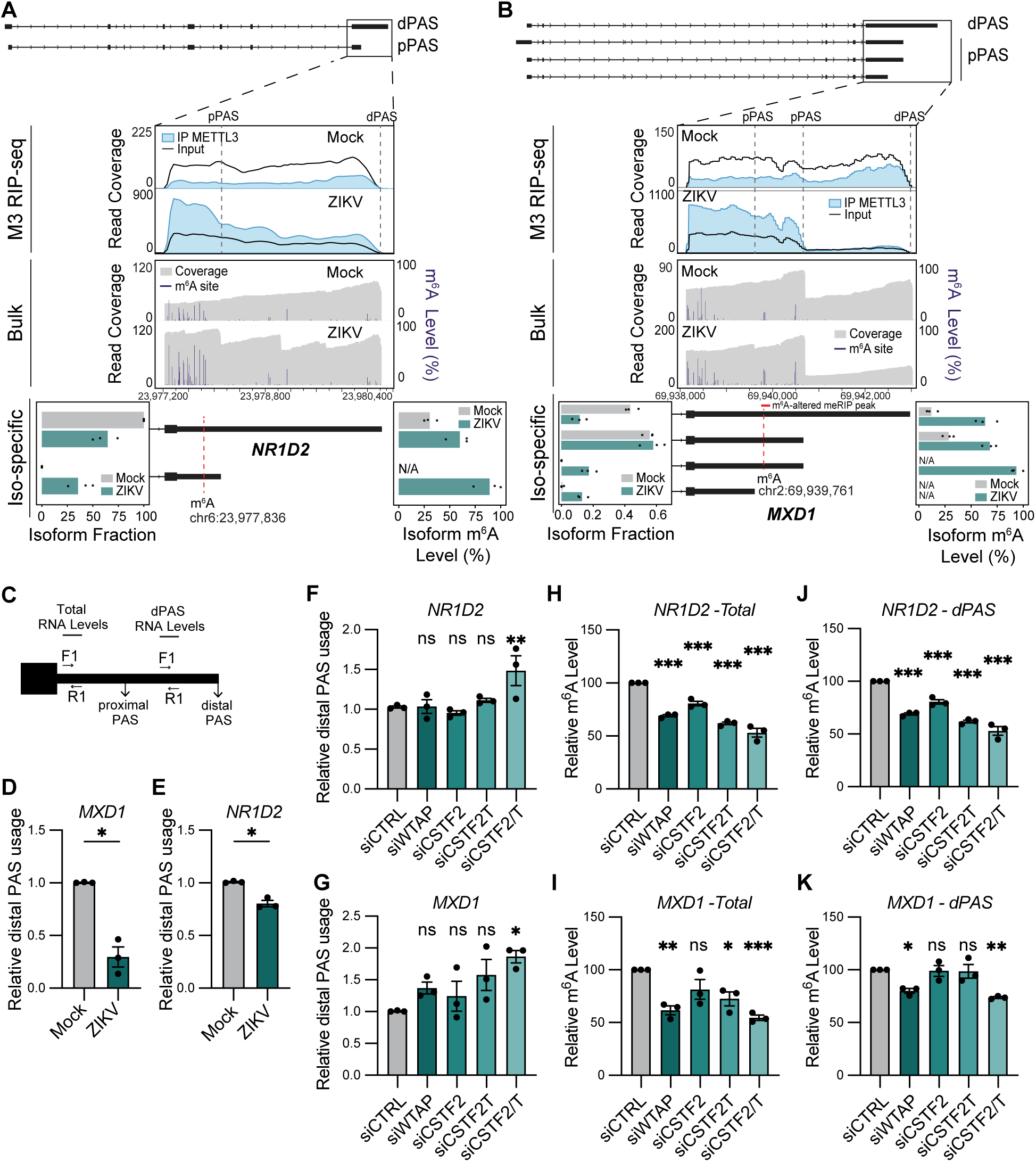
CSTF2/T regulate m^6^A modification of *NR1D2* and *MXD1* independent of their role in alternative polyadenylation. **A)** Representative gene track plot of METTL3 binding and m^6^A methylation in *NR1D2*, along with quantification of isoform fractions and isoform-specific m^6^A levels at the specified site, as determined by Modulator. Distal poly(A) sites (dPAS) and proximal poly(A) sites (pPAS) are labeled by dotted lines in gene track. **B)** Representative gene track plot of METTL3 binding and m^6^A methylation in *MXD1*, along with quantification of isoform fractions and isoform-specific m^6^A levels at the specified site, as determined by Modulator. dPAS and pPAS are labeled by dotted lines in gene track. **C)** Schematic of primers designed to quantify relative dPAS usage by quantifying dPAS RNA levels relative to total RNA levels. **D)** and **E)** RT-qPCR analysis of relative dPAS usage in *MXD1* **(D)** and *NR1D2* **(E)** in Huh7 cells infected with ZIKV (48 hpi, MOI 0.5). **F)** and **G)** RT-qPCR analysis of relative dPAS usage of *NR1D2* **(F)** and *MXD1* **(G)** in Huh7 cells treated with the indicated siRNA followed by ZIKV infection (48 hpi, MOI 0.5). **H)** and **I)** meRIP-RT-qPCR analysis of relative m^6^A level of *NR1D2-Total* **(H)** and *MXD1-Total* **(I)** in Huh7 cells treated with the indicated siRNA followed by ZIKV infection (48 hpi, MOI 0.5). **J)** and **K)** meRIP-RT-qPCR analysis of relative m^6^A level of *NR1D2-dPAS* **(J)** and *MXD1-dPAS* **(K)** in Huh7 cells treated with the indicated siRNA followed by ZIKV infection (48 hpi, MOI 0.5). For all panels, n = 3 biologically independent experiments, with bars indicating mean and error bars showing the standard error. *p < 0.05, **p < 0.01, ***p < 0.001, or not significant (ns) as determined by Welch’s t-test (**D** and **E**) and one-way analysis of variance (ANOVA) with Dunnett’s multiple comparison test (**F**-**K**).

Isoforms that were preferentially m^6^A-modified and targeted by METTL3 were strongly enriched for those generated by tandem 3′UTR APA (**Figure 3E**). Given the strong enrichment of METTL3 near the proximal PAS, we wanted to compare the methylation levels of isoforms utilizing proximal-poly(A) sites (pPAS) and isoforms utilizing distal-poly(A) sites (dPAS). For both *NR1D2* and *MXD1,* the pPAS isoforms had higher m^6^A levels than the dPAS isoforms, mirroring the METTL3 binding preference (**Figure 5A-B**). However, unlike *ALCAM* and *NHSL1*, both pPAS and dPAS isoforms gained m^6^A upon ZIKV infection, suggesting a cleavage-independent mechanism of m^6^A dynamics. Indeed, EJC-mediated suppression of m^6^A acts on short internal exons, like those in *ALCAM* and *NHSL1*, but it cannot account for m^6^A dynamics in long terminal exons, which lie outside the range of influence of the EJC.

Given that CSTF2/T were responsible for IPA-driven m^6^A acquisition on *ALCAM* and *NHSL1*, we next tested whether CSTF2/T also regulate the tandem 3′UTR APA and m^6^A modification of *NR1D2* and *MXD1*. To distinguish dPAS transcripts from total transcript levels, we used two RT-qPCR primer pairs: one upstream of the pPAS to capture all isoforms, and a second downstream of the pPAS to specifically detect transcripts that read through to the dPAS (**Figure 5C**). Using these primer pairs, we confirmed that ZIKV infection decreased dPAS usage of *MXD1* and *NR1D2* (**Figure 5D-E)**. We next performed siRNA knockdown of CSTF2 and/or CSTF2T, followed by RT-qPCR and meRIP-RT-qPCR, in parallel with the experiments performed on *ALCAM* and *NHSL1* (**Figure 5F-K**). Depletion of WTAP did not affect dPAS usage in *NR1D2* or *MXD1* (**Figure 5F-G**), indicating that m^6^A deposition changes do not drive the observed APA shifts. In contrast, combined depletion of CSTF2 and CSTF2T significantly increased dPAS usage, while individual knockdown of either factor had no effect, consistent with their known paralog redundancy in cleavage site selection (**Figure 5F-G**). Strikingly, the requirements for CSTF2/T-mediated regulation of m^6^A on these transcripts were distinct from their requirements for regulating APA. This was most clearly observed for *NR1D2*, where combined CSTF2/T depletion was required to significantly alter dPAS usage of *NR1D2* (**Figure 5F**), but individual depletion of either CSTF2 or CSTF2T was sufficient to reduce m^6^A levels on both total and dPAS *NR1D2* transcripts (**Figure 5H**; **Figure 5J**). Importantly, the dPAS isoform does not undergo CSTF2/T-mediated pPAS cleavage, indicating that the reduction in m^6^A cannot be explained solely by altered pPAS selection. Instead, these findings support an APA-independent role for CSTF2/T in promoting m^6^A deposition on *NR1D2* transcripts. For *MXD1*, the relationship between APA and m^6^A regulation was less clearly dissociated. Combined CSTF2/T depletion was required to both increase dPAS usage and reduce m^6^A levels (**Figure 5G**; **Figure 5I**). Critically, however, this requirement also applied to m^6^A on the dPAS isoform itself (**Figure 5K)**; because this isoform has not undergone pPAS cleavage, its change in m^6^A cannot be a downstream consequence of APA. Together, these findings reveal an APA-independent function for CSTF2/T in promoting m^6^A deposition, distinct from their canonical roles in directing pPAS usage.

### CSTF2 and CSTF2T are required for METTL3 recruitment to target transcripts

Given our finding that CSTF2 and CSTF2T regulate m^6^A on *NR1D2* and *MXD1* independently of cleavage, we next sought to define the mechanism by which CSTF2/T promote methylation at these transcripts. A prior study reported that CSTF2 can stabilize METTL3 interaction with RNAPII^33^, raising the possibility that CSTF2/T promote METTL3 recruitment to our target transcripts. We therefore performed siRNA knockdown of CSTF2 and CSTF2T followed by METTL3 RIP-RT-qPCR (**Figure 6A**). We first examined *ALCAM* and *NHSL1*, which exhibit isoform-specific m^6^A regulation through IPA (**Figure 4**). As expected given their IPA-dependent regulation, combined depletion of CSTF2 and CSTF2T significantly decreased METTL3 binding to *ALCAM* and *NHSL1*, whereas individual knockdown of either factor had minimal effect (**Figure 6B–C**). Depletion of CSTF2 and/or CSTF2T did not affect METTL3 binding to *ACTB* (**Figure 6D**), which does not undergo altered m^6^A or APA during ZIKV infection. In contrast to *ALCAM* and *NHSL1*, METTL3 binding to both total and dPAS transcripts of *NR1D2* and *MXD1* was reduced by individual knockdown of either CSTF2 or CSTF2T (**Figure 6E–H**), confirming the cleavage-independent pattern of m^6^A regulation we observed for these transcripts (**Figure 5**). Together, these results indicate that CSTF2/T promote METTL3 recruitment to all four target transcripts, and that the redundancy requirements for METTL3 RNA binding parallel those for m^6^A regulation: combined CSTF2/T activity is required at IPA targets, while either factor alone is sufficient at cleavage-independent targets.

**Figure 6.**
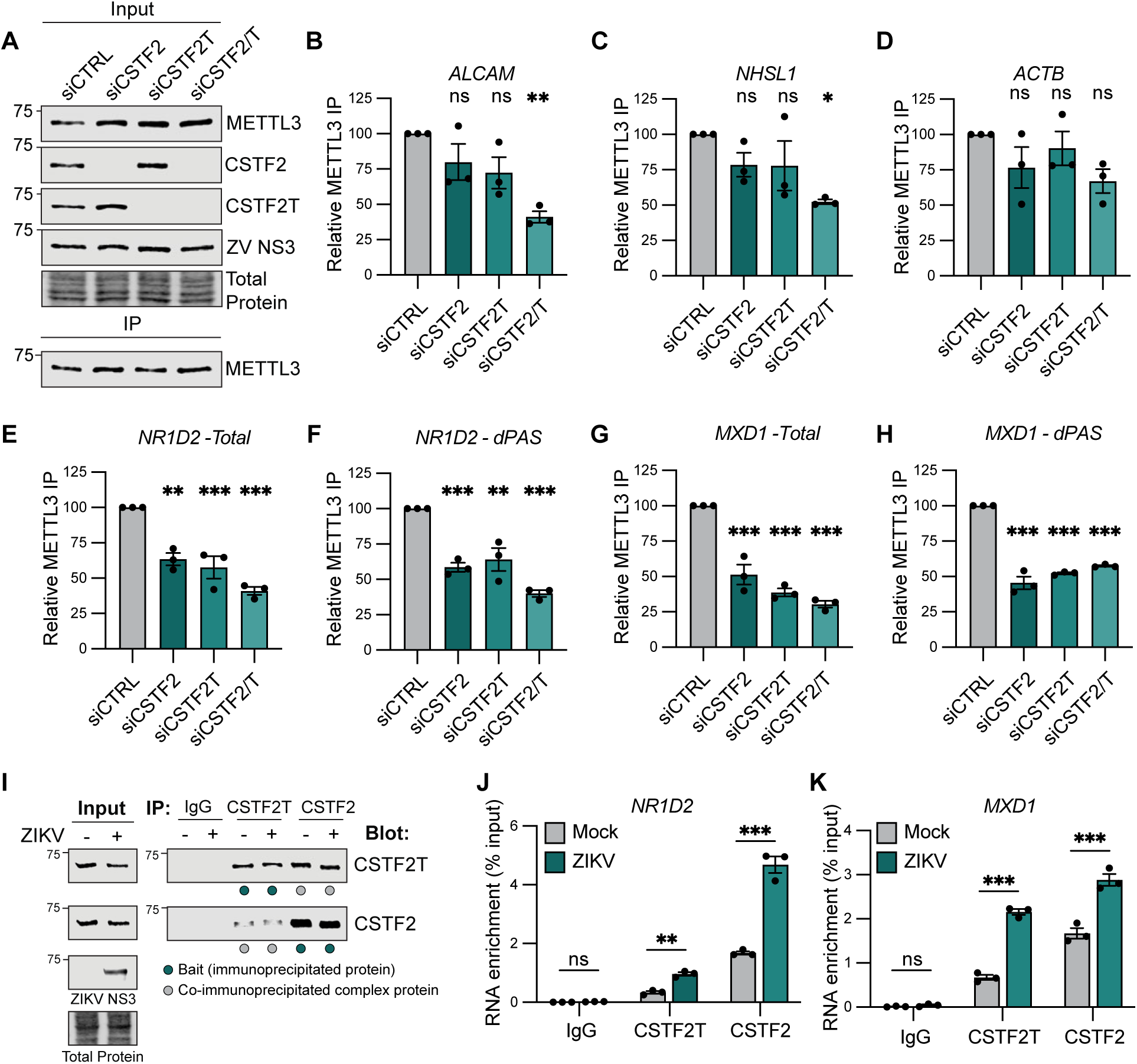
CSTF2 and CSTF2T regulate METTL3 targeting to *NR1D2* and *MXD1* during ZIKV infection. **A)** Representative immunoblots of inputs and immunoprecipitated lysates from Huh7 cells treated with the indicated siRNA followed by ZIKV infection (48 hpi, MOI 0.5), n=3 biologically independent experiments. **B), C),** and **D)** METTL3 RIP-RT-qPCR analysis of relative m^6^A level of *ALCAM* **(B)**, *NHSL1* **(C),** and *ACTB* **(D)** in Huh7 cells treated with the indicated siRNA followed by ZIKV infection (48 hpi, MOI 0.5). **E)** and **F)** METTL3 RIP-RT-qPCR analysis of relative m^6^A level of *NR1D2-Total* **(E)** and *NR1D2-dPAS* **(F)** in Huh7 cells treated with the indicated siRNA followed by ZIKV infection (48 hpi, MOI 0.5). **G)** and **H)** METTL3 RIP-RT-qPCR analysis of relative m^6^A level of *MXD1-Total* **(G)** and *MXD1- dPAS* **(H)** in Huh7 cells treated with the indicated siRNA followed by ZIKV infection (48 hpi, MOI 0.5). **I)** Representative immunoblots of inputs and immunoprecipitated lysates from mock- or ZIKV-infected Huh7 cells (48 hpi, MOI 0.5), n=3 biologically independent experiments. **J)** and **K)** CSTF2/T RIP-RT-qPCR analysis of *NR1D2* **(J)** and *MXD1* **(K)** in Huh7 cells infected with mock or ZIKV (48 hpi, MOI 0.5). For all panels, n = 3 biologically independent experiments, with bars indicating mean and error bars showing the standard error. *p < 0.05, **p < 0.01, ***p < 0.001, or not significant (ns) as determined by one-way analysis of variance (ANOVA) with Dunnett’s multiple comparison test (**B-H**) and two-way ANOVA followed by Šidák’s multiple comparison test (**J**-**K**).

To test whether this CSTF2/T-METTL3 axis is dynamically engaged during infection, we asked whether ZIKV infection increases CSTF2 and CSTF2T binding to *NR1D2* and *MXD1*. We performed CSTF2 and CSTF2T RIP-RT-qPCR in mock- and ZIKV-infected cells and found that ZIKV infection increased both CSTF2 and CSTF2T binding to *NR1D2* and *MXD1* (**Figure 6I–K).** This infection-induced increase in CSTF2/T occupancy thus provides a likely mechanism for the increased METTL3 recruitment and m^6^A acquisition we observe on these transcripts during ZIKV infection.

### m^6^A is enriched upstream of proximal poly(A) sites independently of cleavage, reframing the canonical stop-codon enrichment

Gene-track analyses of *NR1D2* and *MXD1* (**Figure 5A-B**) suggested that METTL3 binding and m^6^A methylation were concentrated upstream of the proximal PAS, a region shared by both pPAS and dPAS isoforms. This raises a broader question: is preferential m^6^A deposition upstream of proximal PAS sites a transcriptome-wide architectural feature, or a gene-specific phenomenon? To assess this, we mapped m^6^A site distribution relative to alternative poly(A) sites in genes with multiple annotated poly(A) sites. m^6^A sites were strongly enriched and highly methylated immediately upstream of PAS1, with a smaller secondary peak upstream of PAS2 and little-to-no density at PAS3 (**Figure 7A-B**). This enrichment and m^6^A methylation level pattern was essentially identical in ZIKV-infected cells (**Figure 7A-B**), indicating that PAS1-proximal m^6^A enrichment is a stable architectural feature of multi-PAS transcripts rather than an infection-induced phenomenon. Furthermore, in genes exhibiting ZIKV-induced APA, defined as those using only one poly(A) site in mock and multiple poly(A) sites in ZIKV, we observed similar enrichment of m^6^A upstream of the PAS1 in both conditions (**Figure 7C**), indicating that this PAS1-proximal enrichment is not a direct consequence of cleavage at the proximal site. Together, these data identify proximal PAS positioning as a transcriptome-wide architectural feature of m^6^A deposition that operates independently of cleavage at the proximal site, raising the possibility that factors influencing PAS1-proximal METTL3 recruitment may be major determinants of m^6^A deposition. Consistent with this possibility, the transcriptome-wide distribution of CSTF2 and CSTF2T binding sites from the POSTAR3 database^70^, which integrates CLIP-seq datasets across diverse cell types and experimental systems, closely resembled the distribution of m^6^A, with strong enrichment upstream of the first proximal PAS, the same region where we observed METTL3 binding (**Figure 7D**).

**Figure 7.**
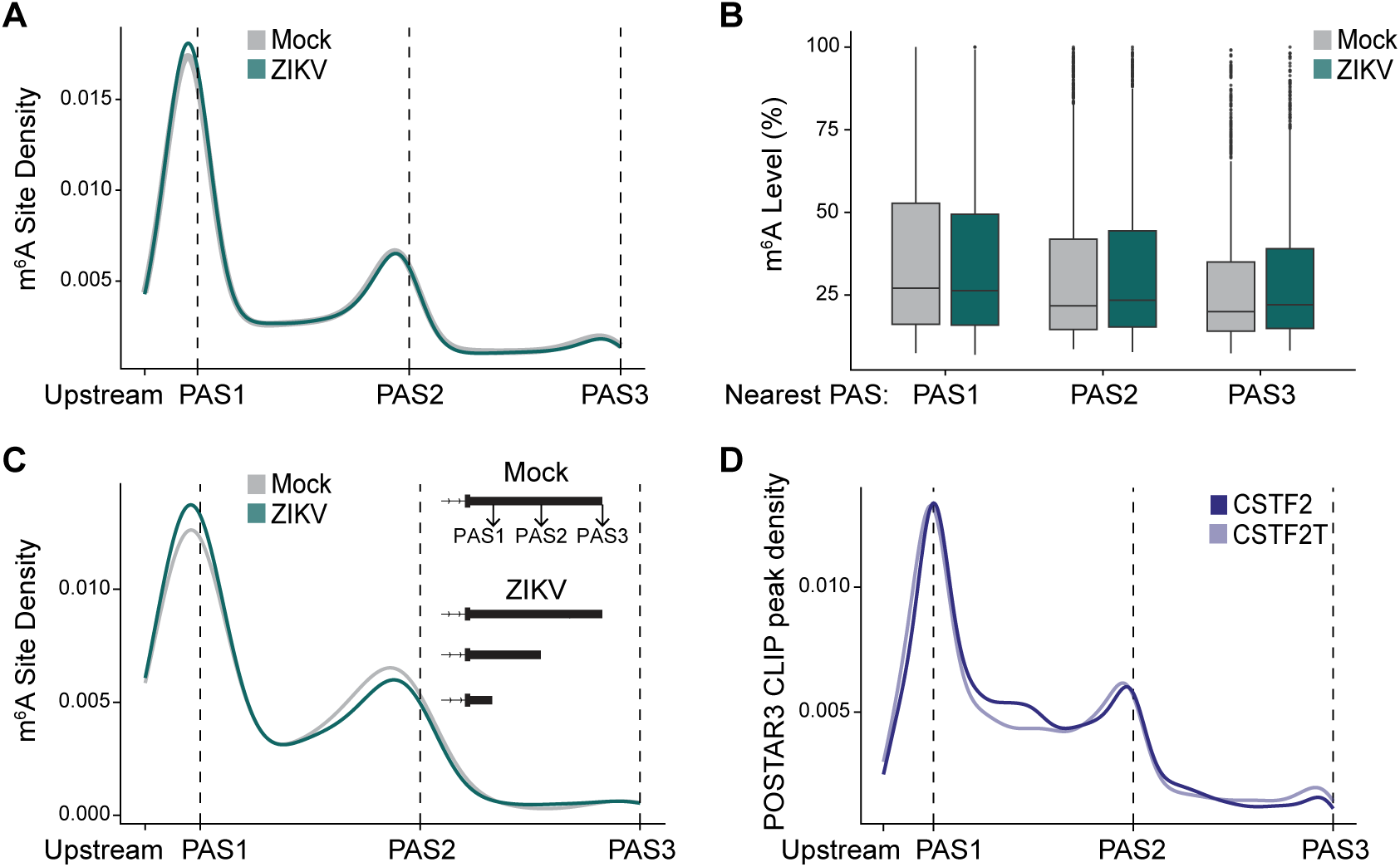
m^6^A is preferentially localized and highly methylated near proximal PAS sites. **A)** Metagene profiles showing the distribution of m^6^A sites across alternative poly(A) sites, as determined by Modulator, in mock- and ZIKV-infected Huh7 cells (48 hpi, MOI 1). **B)** Boxplots showing distribution of methylation level of m^6^A sites that are nearest to the PAS1, PAS2, or PAS3 in mock- and ZIKV-infected Huh7 cells (48 hpi, MOI 1). **C)** Metagene profiles showing the distribution of m^6^A sites across alternative poly(A) sites for genes with ZIKV-induced APA. **D)** Metagene profiles showing the distribution of CSTF2 and CSTF2T POSTAR3^70^ peaks across alternative poly(A) sites determined by Modulator.

## Discussion

This study establishes transcript architecture shaped by alternative polyadenylation as a key determinant of m^6^A dynamics and also defines two mechanistically distinct routes by which the cleavage stimulation factors CSTF2 and CSTF2T direct METTL3 to specific host transcripts during viral infection (**Figure 8**). By integrating GLORI-seq, transcript-level METTL3 RIP-seq, and nanopore DRS, we resolved isoform-level m^6^A regulation that would otherwise be obscured by conventional gene-level approaches. Through this integration, we found that ZIKV-induced m^6^A remodeling occurs by two architectural routes: (i) CSTF2/T-driven IPA, which converts internal exons into terminal exons and thereby exposes previously masked DRACH motifs to METTL3; and (ii) CSTF2/T-mediated METTL3 recruitment near proximal PAS, which operates independently of cleavage at those sites. These data identify CSTF2 and CSTF2T as central regulators of ZIKV-induced remodeling of the host m^6^A mRNA landscape^22^.

**Figure 8.**
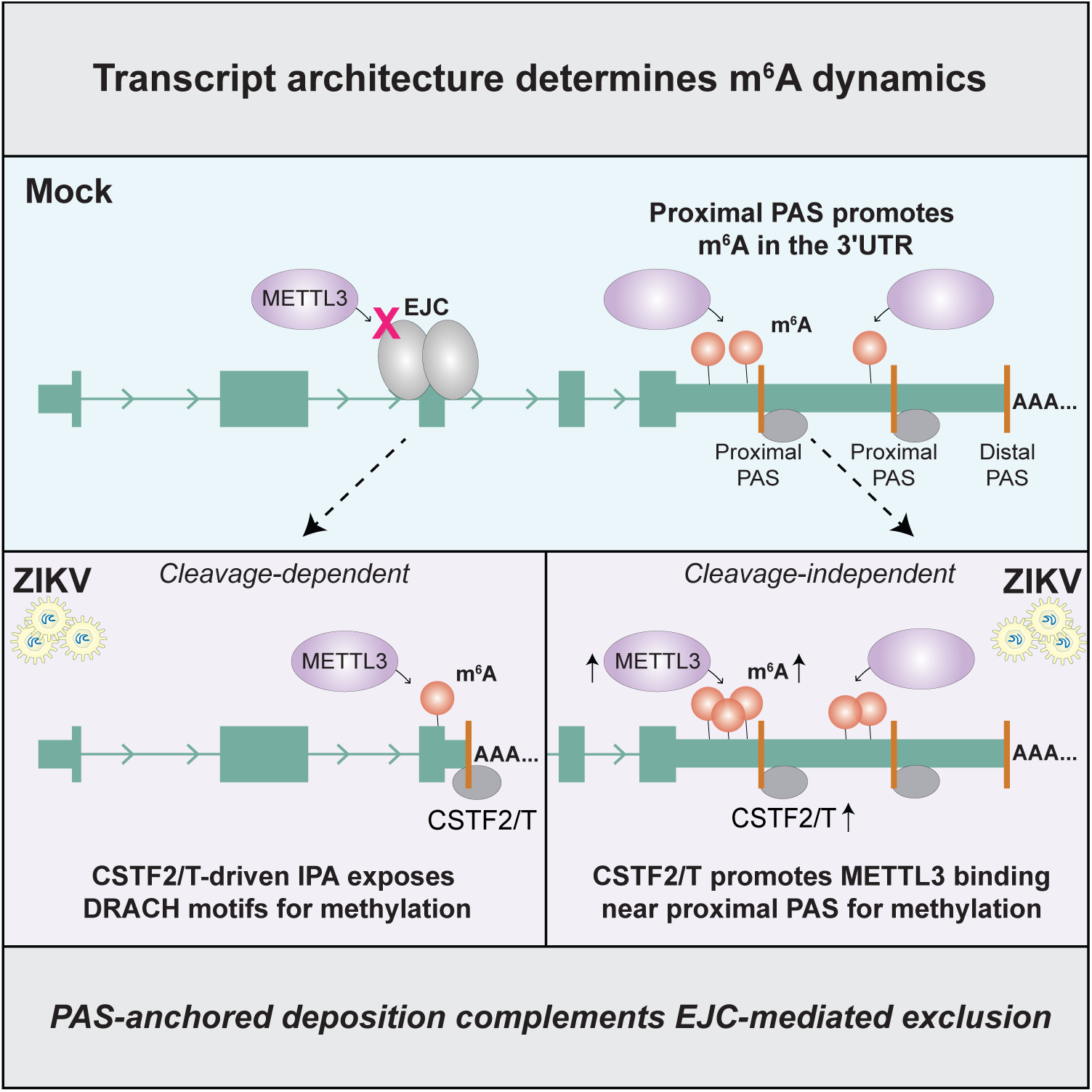
**Transcript architecture shaped by alternative polyadenylation determines where m^6^A accumulates, with m^6^A enriched upstream of proximal polyadenylation sites**. During Zika virus infection, the cleavage stimulation factors CSTF2 and CSTF2T regulate METTL3 targeting by two routes: intronic polyadenylation that exposes new DRACH motifs in terminal exons, and recruitment of METTL3 near proximal polyadenylation sites independent of cleavage.

Perturbing CSTF2 and CSTF2T individually and in combination revealed two separable mechanisms of METTL3 targeting. At IPA targets such as *ALCAM* and *NHSL1*, CSTF2 and CSTF2T act redundantly to drive cleavage at the proximal PAS, and combined depletion is required to reverse both the isoform switch and the m^6^A gain. The same genomic adenosine is methylated only when ZIKV-induced IPA unmasks it as a terminal-exon site. Recent work established that splice-junction-proximal DRACH motifs are occluded by the EJC and by broader exon-intron boundary inhibition, restricting METTL3-mediated m^6^A deposition to central regions of long internal exons and terminal exons^23–26^. Mechanistically, IPA converts an internal exon into a terminal exon, removing the downstream exon–exon junction that would otherwise limit METTL3 access, thereby exposing newly terminal DRACH motifs for methylation. m^6^A acquisition is therefore not an incidental byproduct of altered RNA processing but rather a direct architectural consequence of removing an exon-exon junction. Together, these findings indicate that the m^6^A landscape is not static, but can be dynamically rewritten by IPA shifts, extending the exon-junction-exclusion model of m^6^A topology. In this framework, IPA acts as a regulatory input into m^6^A deposition by altering which transcript regions are permissive for methylation as PAS choice changes across cellular states.

In contrast to their redundant requirement at IPA targets, CSTF2 and CSTF2T function non-redundantly to recruit METTL3 at cleavage-independent targets. For *NR1D2* and *MXD1*, depletion of CSTF2/T reduces METTL3 binding and m^6^A on all isoforms including the dPAS isoform, suggesting APA-independent regulation. Prior work in pancreatic cancer established that CSTF2 slows RNAPII elongation and thereby promotes co-transcriptional METTL3 recruitment^33^. Our data extend this co-transcriptional model by revealing the paralog CSTF2T as an additional regulator of METTL3 recruitment, by showing that this model is dynamically engaged in response to viral infection, and by linking CSTF2/T to m^6^A concentration near proximal PAS sites. The co-occupancy of m^6^A, METTL3, and CSTF2/T immediately upstream of proximal poly(A) sites (**Figure 5A-B**; **Figure 7**), together with ZIKV-induced increases in CSTF2 and CSTF2T binding to target transcripts (**Figure 6J-K**), supports a positional mechanism in which cleavage-factor binding itself, rather than the cleavage reaction, promotes METTL3 binding to local DRACH motifs. As shown in pancreatic cancer^33^, this mechanism may be driven by RNAPII elongation dynamics, where CSTF2/T-mediated slowing of RNAPII elongation stabilizes METTL3 binding on nascent transcripts, promoting m^6^A modification. Notably, another member of the m^6^A methyltransferase complex, VIRMA, has been associated with 3′-end processing machinery^71^, providing a plausible model within which CSTF2/T-mediated METTL3 recruitment could occur. Whether CSTF2/T and METTL3 directly associate in a complex on these transcripts during infection remains to be established.

Our transcriptome-wide analyses of APA and m^6^A show that m^6^A is enriched immediately upstream of PAS1, with a smaller, sequential enrichment upstream of PAS2 across multi-PAS genes, and this pattern is preserved between mock and ZIKV-infected conditions (**Figure 7**). The canonical “stop-codon-and-3′UTR” distribution of m^6^A^10, 11^ is therefore more accurately described as a “proximal-PAS-anchored m^6^A enrichment”, with consequences that generalize well beyond infection models. Current literature on m^6^A has emphasized cis-acting sequence context, which can hard code m^6^A stoichiometry at the level of individual sites^72^, and exon-junction-mediated suppression, in which the EJC excludes METTL3 within ∼150 nt of exon-exon junctions^23–25^, with broader exon-intron boundary inhibition focuses m^6^A onto terminal exons^26^. While this mechanism explains why m^6^A modification preferentially occurs in terminal exons, it does not specify where within the 3′UTR m^6^A accumulates. Our data suggest that proximal-PAS positioning acts as a positive determinant: m^6^A is enriched immediately upstream of proximal poly(A) sites across the transcriptome, and CSTF2 and CSTF2T are a likely route through which METTL3 is recruited to these regions, though the factors that govern PAS-anchored m^6^A enrichment more broadly remain to be defined. EJC suppression and PAS-anchored deposition therefore work in opposite directions, with one specifying where m^6^A is *excluded* near splice junctions, and the other specifying where it *accumulates* within the 3′UTR.

Together, our findings demonstrate that ZIKV reshapes host m^6^A on a defined subset of transcripts by driving CSTF2/T-dependent IPA and by recruiting METTL3 near proximal PAS sites. Importantly, this occurs without changes in the expression or localization of the core m^6^A machinery^22^. Other viruses also alter host m^6^A but through distinct routes, including retargeting of METTL3 by viral proteins or host RBPs to specific cellular transcripts^42, 43^. The mechanism we describe is distinct in two ways: ZIKV co-opts a host 3′-end-processing factor rather than a viral effector, and, at a subset of targets, CSTF2/T act by reshaping transcript architecture itself rather than by directing METTL3 to pre-existing transcripts. Given that several viruses (vesicular stomatitis virus, influenza A virus, and herpes simplex virus 1) also induce global 3′UTR shortening during infection^35, 73–76^, CSTF2/T-mediated retargeting of METTL3 may operate alongside 3’-end-processing changes previously described for those viruses, thereby altering m^6^A distribution. Consistent with a broader *Flaviviridae*-level mechanism, the short-isoform induction observed for *ALCAM* and *NHSL1* during ZIKV infection also occurs in DENV and WNV infection (**Figure S5**), pointing to an induced signal upstream of CSTF2/T. During infection, APA can be driven by altered 3′-end-processing factor activity and changes in RNAPII elongation kinetics^35, 76^, but the specific upstream signaling pathways engaged during ZIKV infection remain to be defined. m^6^A remodeling is well positioned to rapidly respond to infection given its known roles in RNA stability, translation, and innate immune signaling^4, 21^. ZIKV antagonizes interferon signaling by hijacking the CUL3-RING E3 ubiquitin ligase complex to target and degrade STAT2^47, 48^. Notably, *STAT2* and multiple substrate adaptors for CUL3-RING E3 ubiquitin ligases, including *KCTD5* and *KCTD10*, gain m^6^A upon ZIKV infection. This raises the possibility that m^6^A remodeling of these transcripts represents an additional regulatory layer that tunes antiviral defenses at the RNA level, although whether this reflects a host countermeasure or viral co-option, or whether infection-induced m^6^A changes on specific transcripts are functionally consequential for infection outcome, is unknown and will require site-directed disruption of the specific m^6^A modifications identified here, as well as additional characterization.

Although most m^6^A-modified genes are METTL3-bound in our data, ∼20% of m^6^A-altered genes showed concordant changes in METTL3 binding by transcript-level RIP-seq (**Figure S2D**), pointing to additional regulatory layers. One possibility is that infection alters METTL3 catalytic activity through post-translational modifications. Indeed, multiple post-translational modifications, including SUMOylation, phosphorylation, ubiquitination, and acetylation, have been shown to regulate METTL3 catalytic activity^77^, and viral infection has been reported to rewire these post-translational modifications to modulate m^6^A^21, 78^. Altered targeting or activity of the m^6^A demethylases ALKBH5 and FTO may also contribute^44, 79–83^. Defining how other m^6^A machinery proteins are coordinately regulated during ZIKV infection will reveal additional regulators of infection-associated m^6^A changes.

Integrating GLORI-seq, transcript-resolved DRS, and METTL3 RIP-seq was essential to this work. Gene-level m^6^A increases identified by GLORI-seq frequently reflected infection-induced shifts in isoform usage detectable only by DRS: 53% of GLORI-detected m^6^A-increased genes contained isoform-specific m^6^A patterns by DRS (**Figure 3I**), and a substantial fraction of ZIKV-induced METTL3 binding changes were detected only at the transcript level and missed by gene-level analysis (**Figure 2D; Figure S2E**). This illustrates how single-method approaches can mistake compositional shifts in RNA architecture for uniform changes in modification per site. These findings underscore the value of orthogonal m^6^A profiling approaches for uncovering layers of epitranscriptomic regulation that are otherwise obscured.

In summary, this work provides a single-nucleotide, isoform-specific map of m^6^A dynamics during ZIKV infection, enabling future functional studies on how differentially methylated sites affect splicing, translation, and stability. More broadly, the principles described here establish that transcript architecture sets the m^6^A landscape, that cleavage factors recruit METTL3 to defined positions, and that APA actively rewrites this architecture. These principles should apply in any context where APA is dynamically regulated, including development, cancer, and cellular stress.

## Acknowledgements

We thank current and past members of the Horner Lab for valuable feedback and discussion; the Duke Functional Genomics Core Facility; and the Weill Cornell Genomics Resources Core Facility. Research on m^6^A in the Horner lab has been supported by Burroughs Wellcome Fund and National Institutes of Health (NIH) grant R01AI125416. Additionally, this material is based upon work supported by the National Science Foundation Graduate Research Fellowship (C.J.A), and the CMB training grant T32GM142605 (C.J.A). C.E.M. was supported by the Katherine and John Bleckman Foundation, and the NIH (U54AG089334, R01AI125416). T.M.N. was supported by a Medical Scientist Training Program grant from the National Institute of General Medical Sciences of the National Institutes of Health under award number: T32GM152349 to the Weill Cornell/Rockefeller/Sloan Kettering Tri-Institutional MD-PhD Program. T.M.N. was also supported with computational resources from the National Science Foundation ACCESS Allocation Request BIO240371. Additional computations by T.M.N. in this paper were run with HPC resources supported by the Scientific Computing Unit at Weill Cornell Medicine. E.M.V. was supported by the Weill Cornell ACCESS Summer Internship Program. K.D.M. was supported by the NIH (1RM1HG011563).

## Author Contributions

Conceptualization, C.J.A., T.M.N., C.E.M., and S.M.H.; Methodology, C.J.A., T.M.N., N.H., and S.D.V.; Software, T.M.N., N.H., and S.D.V.; Formal Analysis, C.J.A., T.M.N., N.H., S.D.V., E.L., E.S., M.G., and E.M.V.; Investigation, C.J.A., M.T. P.C., J.P., K.R., and S.A.T.; Writing – Original Draft, C.J.A., and S.M.H.; Writing – Review & Editing, all authors; Visualization, C.J.A. and S.M.H.; Supervision, K.D.M., C.E.M., and S.M.H.; Funding Acquisition, K.D.M., C.E.M., and S.M.H.

## Declaration of Interests

C.E.M. is a co-founder of Biotia.

## Declaration of generative AI and AI-assisted technologies in the writing process

During the preparation of this work, the authors used Perplexity (including its Deep Research and Computer features, with access to Claude Opus 4.7 and 4.8 from Anthropic), and ChatGPT (OpenAI) to improve readability and language and to assist with coding scripts. After using these tools, the authors reviewed and edited the content as needed and take full responsibility for the content of the publication.

## Data and code availability

Raw sequencing data will be deposited upon publication. Relevant code and outputs can be found in this GitHub repository: https://github.com/Theo-Nelson/horner-zika/tree/main.

## Supplementary Figures

**Figure S1: Related to Figure 1.**
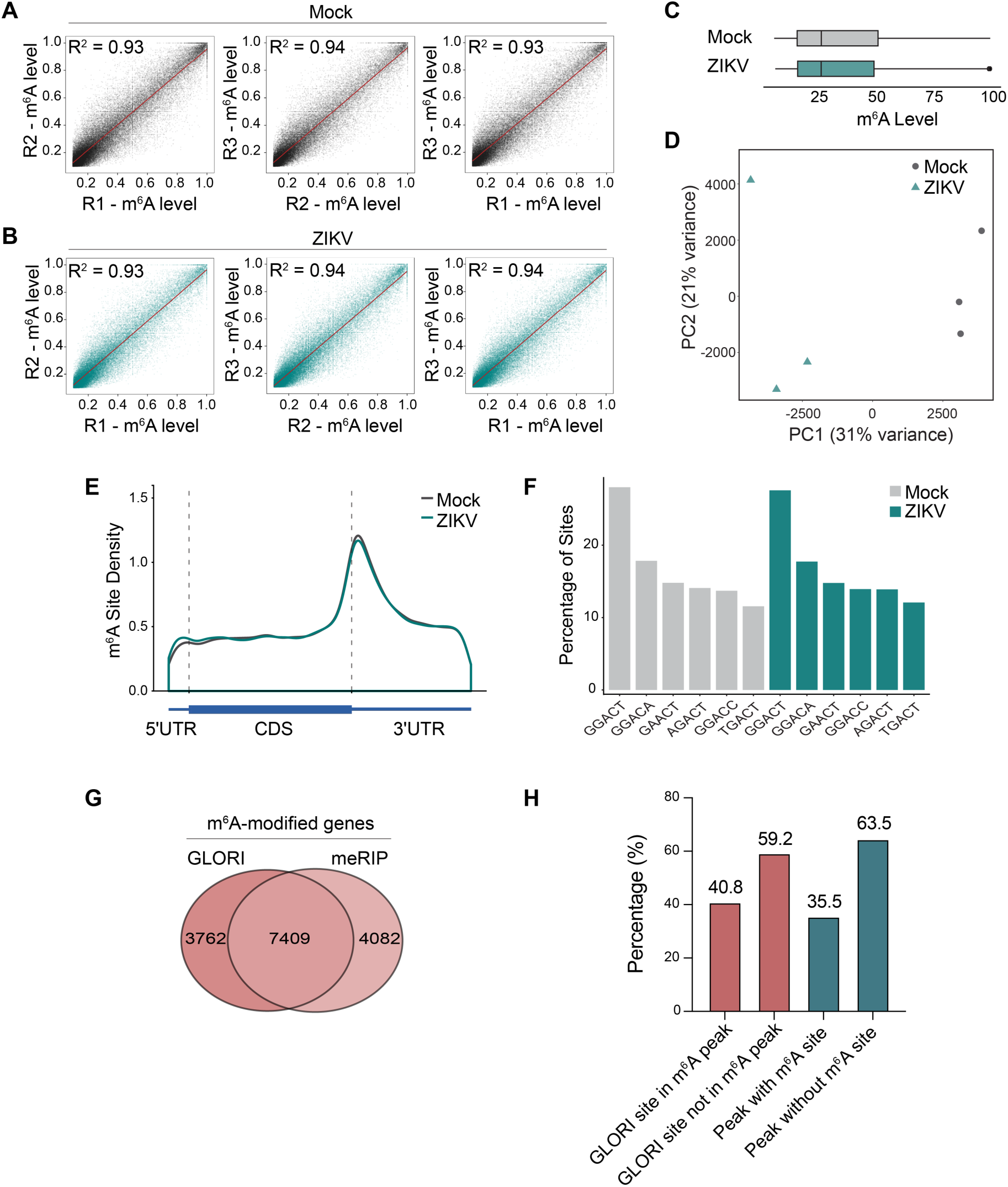
GLORI-seq accurately detects m^6^A methylation of cellular RNAs during ZIKV infection. **A)** and **B)** The correlation of m^6^A levels for sites detected by GLORI-seq between three biologically independent samples of mock- **(A)** or ZIKV-infected **(B)** Huh7 cells (48 hpi, MOI 1). **C)** Boxplots showing distribution of methylation level of m^6^A sites in mock and ZIKV conditions. **D)** Principal component analysis of GLORI-seq m^6^A stoichiometry rates for mock and ZIKV conditions. **E)** Metagene profiles showing the distributions of mock and ZIKV m^6^A sites across the mRNA molecule. **F)** The top 6 motifs across mock and ZIKV m^6^A sites and the proportion of m^6^A sites they contain. **G)** Venn diagram showing the overlap of genes detected to be m^6^A-modified during infection in GLORI-seq and meRIP-seq^22^. **H)** Histogram showing the overlap percentage between GLORI-detected m^6^A sites and meRIP-seq peaks^22^.

**Figure S2: Related to Figure 2.**
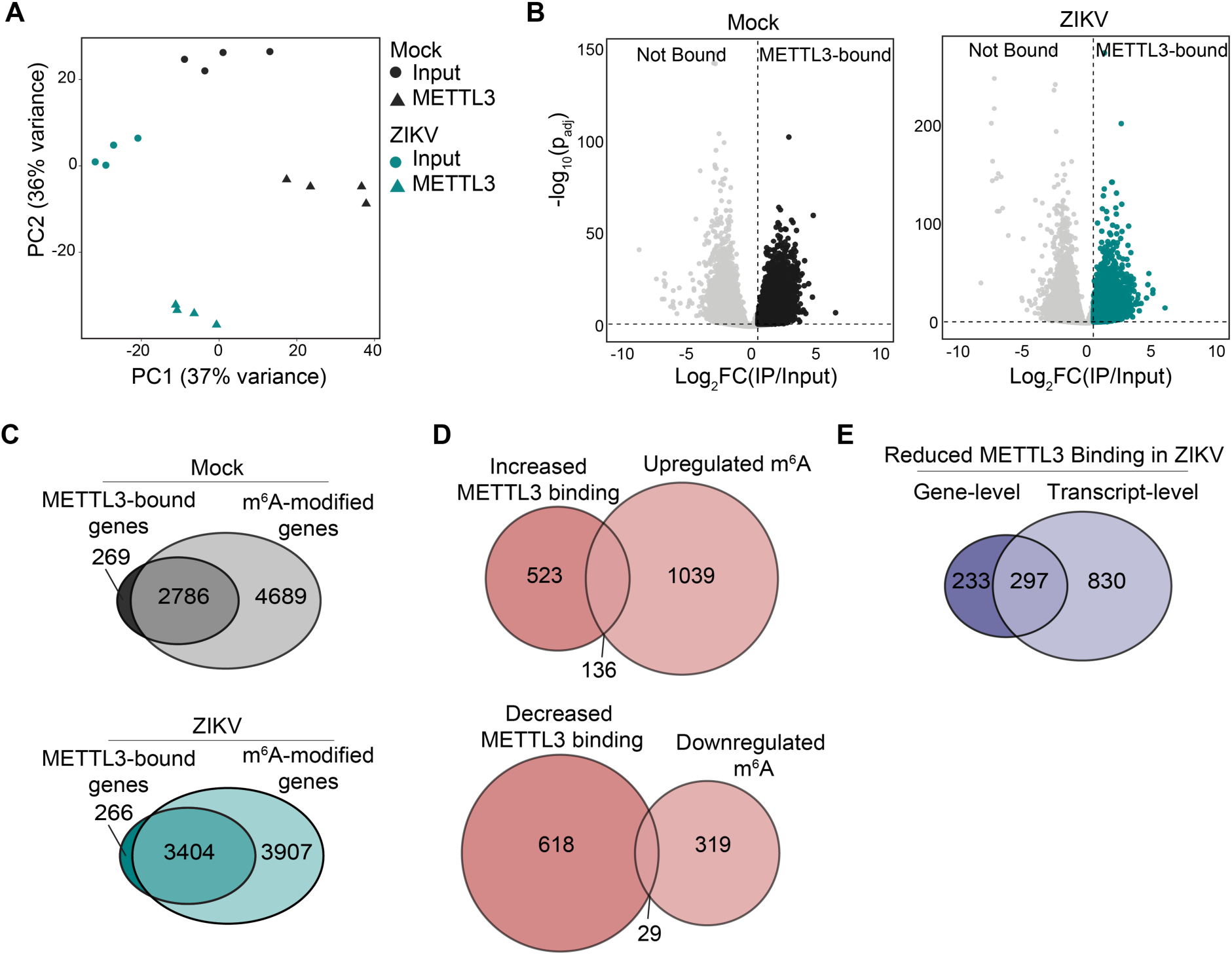
METTL3 RIP-seq captures METTL3-bound cellular RNAs that are m^6^A-modified. **A)** Principal component analysis of variance-stabilized transcript-level counts from METTL3 RIP-seq transcript-level analysis. **B)** Volcano plot showing METTL3-bound genes (IP/Input Log_2_FC ≥ 0.58, p_adj_ < 0.05) in mock- (left) and ZIKV-infected (right) Huh7 cells (48 hpi, MOI 1). Black and teal dots represent METTL3-bound genes in mock and ZIKV conditions, respectively. **C)** Venn diagram comparing genes that are METTL3-bound and genes that are m^6^A-modified in mock (top) and ZIKV (bottom) conditions. **D)** (Top) Venn diagram comparing genes with increased METTL3 binding and genes that contain increased m^6^A. (Bottom) Venn diagram comparing genes with decreased METTL3 binding and genes that contain decreased m^6^A. **E)** Venn diagram showing the overlap of genes detected to have decreased METTL3 binding in the gene-level analysis (STAR-based workflow) and the transcript-level analysis (Salmon-based workflow). For panels **(C)**, **(D)**, and **(E)**, only genes that pass coverage thresholds in both data sets (reads≥15 in GLORI-seq data, reads≥10 in METTL3 RIP-seq data) were overlapped to avoid confounding biological discordance with discrepancies in sequencing coverage.

**Figure S3: Related to Figure 3.**
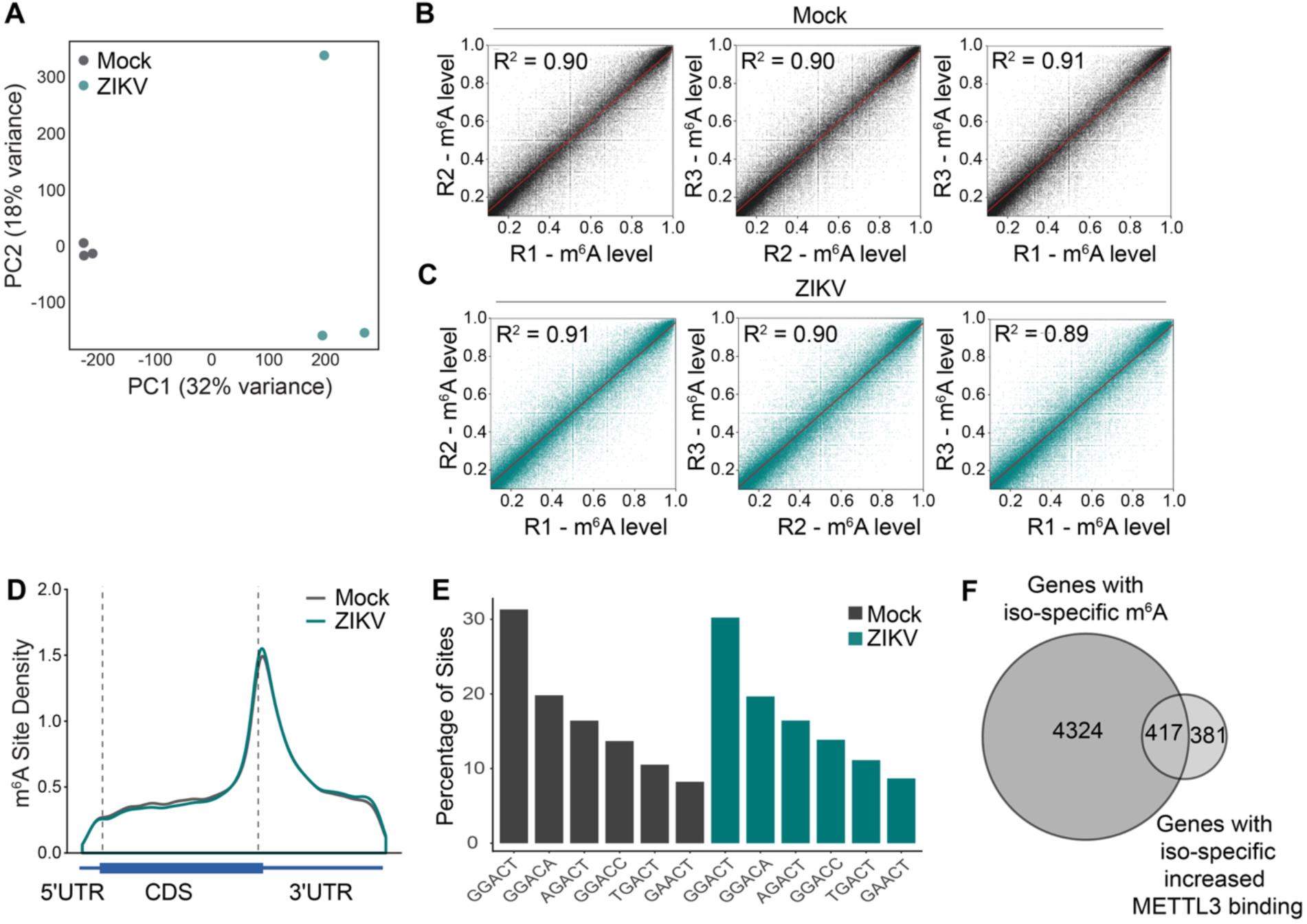
DRS accurately detects m^6^A methylation of cellular RNAs during ZIKV infection. **A)** Principal component analysis of DRS m^6^A stoichiometry rates for mock and ZIKV conditions. **B)** and **C)** The correlation of m^6^A levels for sites detected by DRS between three biologically independent samples of mock- **(B)** or ZIKV-infected **(C)** Huh7 cells (48 hpi, MOI 1). **D)** Metagene profiles showing the distributions of mock and ZIKV m^6^A sites across the mRNA molecule. **E)** The top 6 motifs across mock and ZIKV m^6^A sites and the proportion of m^6^A sites they contain. **F)** Venn diagram comparing genes with iso-specific increased METTL3 binding and genes that contain iso-specific m^6^A.

**Figure S4: Related to Figure 4.**
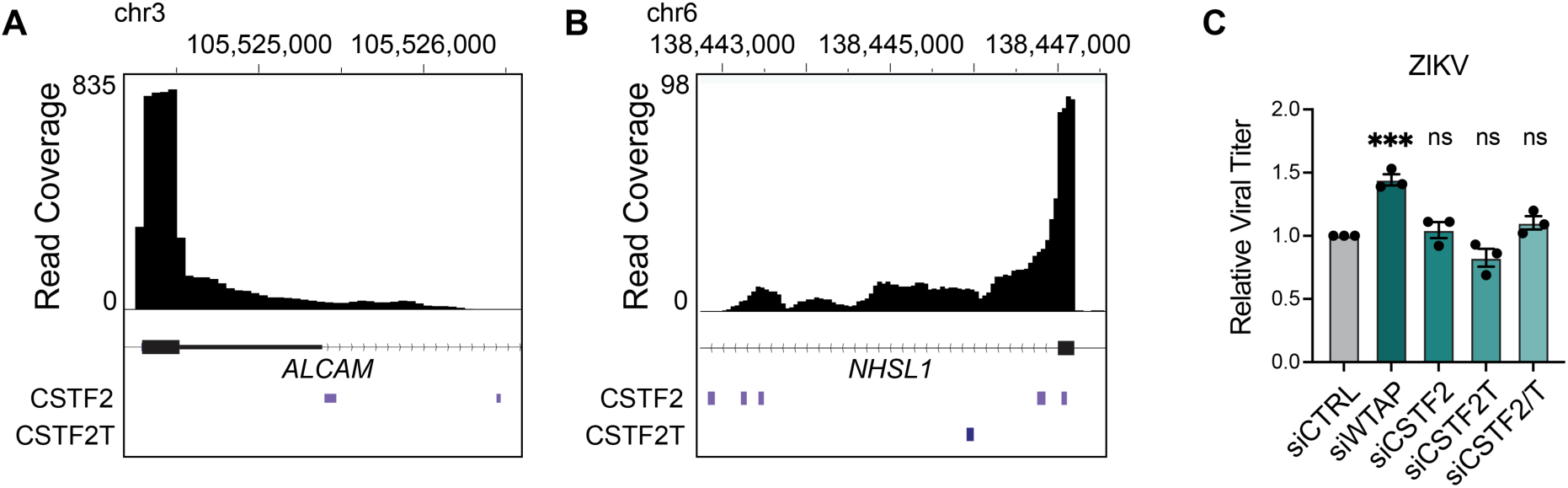
CSTF2/T bind to *ALCAM* and *NHSL1* near IPA events and do not affect ZIKV viral titer. **A)** and **B)** Gene track plots of *ALCAM* and *NHSL1* during ZIKV infection annotated with CSTF2/T ENCODE^61^ eCLIP peaks. **C**) Focus-forming assay of supernatants harvested from Huh7 cells treated with the indicated siRNAs followed by ZIKV infection (48 hpi, MOI 0.5). n = 3 biologically independent experiments, with bars indicating mean and error bars showing the standard error. ***p < 0.001, or not significant (ns) as determined by one-way analysis of variance (ANOVA) with Dunnett’s multiple comparison test.

**Figure S5: Related to Figure 4.**
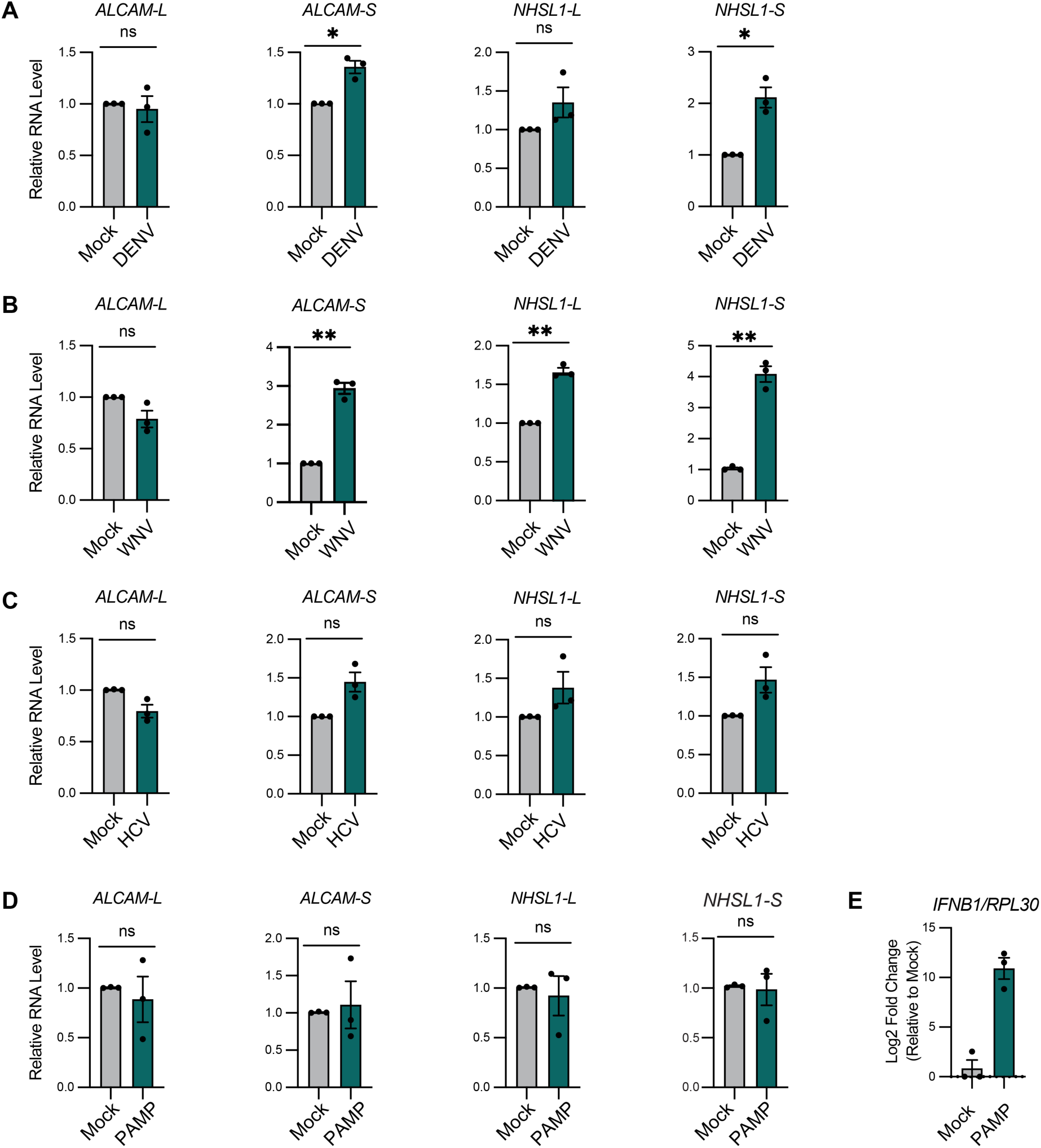
Intronic polyadenylation of *ALCAM* and *NHSL1* upon viral infection is conserved across multiple flaviviruses. A) RT-qPCR analysis (relative to *RPL30*) of RNA expression of *ALCAM-L*, *ALCAM-S*, *NHSL1-L*, *NHSL1-S* upon DENV infection (48 hpi, MOI 1). **B)** RT-qPCR analysis (relative to *RPL30*) of RNA expression of *ALCAM-L*, *ALCAM-S*, *NHSL1-L*, and *NHSL1-S* upon WNV infection (48 hpi, MOI 0.01). **C)** RT-qPCR analysis (relative to RPL30) of RNA expression of *ALCAM-L*, *ALCAM-S*, *NHSL1-L,* and *NHSL1-S* upon HCV infection (48 hpi, MOI 1). **D)** RT-qPCR analysis (relative to *RPL30*) of RNA expression of *ALCAM-L*, *ALCAM-S*, *NHSL1-L,* and *NHSL1-S* upon transfection with HCV PAMP (8 hrs). **E)** RT-qPCR analysis (relative to *RPL30*) of RNA expression of *IFNB1* upon transfection with HCV PAMP (8 hrs). For all panels, n = 3 biologically independent experiments, with bars indicating mean and error bars showing the standard error. *p < 0.05, **p < 0.01, or not significant (ns) as determined by Welch’s t-test.

## Supplemental Note

### Note S1: Related to Figure 1

To assess concordance between GLORI-seq and our previously published meRIP-seq data^22^, we compared genes and sites identified as m^6^A-modified by each method. We restricted comparison to genes that passed read coverage thresholds in both datasets, ensuring that differences in m^6^A detection reflected biological or technical differences between methods rather than insufficient read coverage. We found that 64% of m^6^A-modified genes identified by meRIP-seq were also detected to be m^6^A-modified by GLORI-seq (**Figure S1G**). Of the 31,647 meRIP peaks previously detected during *Flaviviridae* infection, 36% directly overlapped one or more GLORI-detected m^6^A sites, and 41% of GLORI m^6^A sites were within a previously defined meRIP peak (**Figure S1H**). Of the 51 m^6^A-altered genes and 54 m^6^A-altered peaks that we previously identified^22^, 25% of m^6^A-altered genes also had dynamic m^6^A by GLORI-seq, while only 14% of m^6^A-altered peaks directly overlapped dynamic GLORI m^6^A sites. Together, these results indicate that meRIP-seq can reliably detect m^6^A-modified genes but has limited precision in resolving the exact sites and dynamic changes in methylation compared to single-nucleotide-resolution approaches like GLORI-seq.

## METHODS

### Cell Culture

Huh7 and Huh7.5 cells (gifts of Dr. Michael Gale Jr., University of Washington)^84^, and Vero cells (ATCC) were grown in Dulbecco’s modification of Eagle’s medium (DMEM, Mediatech) supplemented with 10% fetal bovine serum (HyClone), 1X minimum essential non-essential amino acids (Thermo Fisher), and 25 mM HEPES (Thermo Fisher), referred to as complete DMEM (cDMEM). C6/36 cells (ATCC: CRL-1660) were grown in Eagle’s minimum essential media (ATCC) supplemented with 10% fetal bovine serum (HyClone), 1 mM Sodium pyruvate (Thermo Fisher), and 1X minimum essential non-essential amino acids (Thermo Fisher). Cells were verified using the GenePrint STR kit (Duke DNA Analysis Facility) and were confirmed mycoplasma free by the MycoStrip Mycoplasma Detection Kit (Invivogen).

### Viruses

ZIKV (ZIKV/*Homo sapiens*/PRI/PRVabc59/2015)^85^, DENV (dengue virus 2 Thailand/NGS-C/1944)^86^, and WNV (WNV strain 3000.0259 isolated in New York in 2000)^87^ stocks were prepared in C6/36 cells and titered on Vero cells using a focus-forming assay, as described previously^88^. Infectious stocks of a cell culture-adapted strain of genotype 2A JFH1 HCV^89^ were generated and titered on Huh7.5 cells by focus-forming assay, as described. For viral infections, cells were incubated in a low volume of serum-free DMEM containing virus at the indicated multiplicity of infection (MOI) for 3 h, followed by replacement of infected media with cDMEM.

### Focus-forming assay for viral titer

Supernatants were harvested from ZIKV-infected cells 48 hpi, serially diluted, and used to infect naïve Vero cells in triplicate wells of a 48-well plate for 3 h before overlay with methyl cellulose (Millipore Sigma, M0512). At 72 hpi, cells were washed with phosphate-buffered saline (PBS) and fixed with 1:1 methanol:acetone. Cells were blocked with 5% milk in PBS with 0.1% Tween (PBS-T) and then immunostained with M-anti-4G2 antibody generated from the D1-4G2-4-15 hybridoma cell line against the flavivirus envelope protein (ATCC; 1:2,000). Infected cells were visualized by incubating with horseradish peroxidase (HRP)-conjugated secondary antibody (1:500) and the VIP Peroxidase Substrate Kit (Vector Laboratories). The titer (focus-forming units per milliliter) was calculated from the average number of 4G2-positive foci at x10 magnification, relative to the amount and dilution of virus used.

### HCV PAMP treatment

HCV PAMP RNA was prepared by *in vitro* transcription, as described previously^67, 90^. 2.5 μg of HCV PAMP RNA was transfected into cells for 8 h using the Mirus mRNA transfection kit.

### Antibodies

For immunoblotting, the following primary antibodies were used: R-anti-METTL3 (Abcam, ab195352, 1:1,000), M-anti-METTL3 (Proteintech, 67733-1-IG, 1:1,000), R-anti-ZIKV NS3 (GeneTex, GTX133320, 1:1,000), R-anti-WTAP (Abcam, ab195380, 1:1,000), R-anti-CSTF2 (Bethyl Labs, A301-487A, 1:1,000), M-anti-CSTF2 (Proteintech, 68520-1-IG, 1:2,000), R-anti-CSTF2T (Bethyl Labs, A301-092A, 1:1,000). For RNA immunoprecipitation, the following antibodies were used: R-anti-METTL3 (Abcam, ab195352, 3μg), R-anti-CSTF2 (Bethyl Labs, A301-487A, 3μg), R-anti-CSTF2T (Bethyl Labs, A301-092A, 3μg), R-anti-IgG (Sigma-Aldrich, PP64B, 3μg).

### siRNA treatment

Cells were transfected with siRNA against indicated targets 24 h before infection using Lipofectamine RNAiMax (Thermo Fisher) according to the manufacturer’s protocol, with media changed 4 h after transfection. Depletion of siRNA targets were confirmed by immunoblot analysis. siRNAs used included siWTAP (Qiagen, SI00069853), siCSTF2 (s3684 and s3686, Ambion), siCSTF2T (s23471 and s23472, Ambion), siCSTF2/T (s3684, s3686, s23471, and s23472, Ambion).

### Immunoblotting

Cells were lysed in a modified radioimmunoprecipitation assay buffer (TX-100-RIPA) (50 mM Tris pH 7.5, 150 mM NaCl, 5 mM EDTA, 0.1% SDS, 0.5% sodium deoxycholate, and 1% Triton X-100) supplemented with protease inhibitor (Sigma-Aldrich) and Halt phosphatase inhibitor (Thermo Fisher) at 1:100, and lysates were clarified by centrifugation. Quantified protein, as determined by Bradford assay (Bio-Rad), was resolved by 10% or 4-20% SDS/polyacrylamide gel electrophoresis (PAGE) and transferred to nitrocellulose membranes in the Trans-Blot Turbo buffer (Bio-Rad) using the Turbo Transfer system (Bio-Rad), followed by staining with Revert Total Protein Stain (Licor Biosciences). Membranes were blocked with 3% bovine serum albumin in PBS-T, probed with primary antibodies against proteins of interest, washed with PBS-T, incubated with species-specific HRP-conjugated secondary antibodies (Jackson ImmunoResearch, 1:5,000) or fluorescent secondary antibodies (Licor Biosciences, 1:5,000), washed again with PBS-T, and treated with Clarity Western ECL substrate (Bio-Rad). Imaging was performed using a LICOR Odyssey FC.

### RT-qPCR

Total cellular RNA was extracted using TRIzol extraction (Thermo Fisher) and diluted to equivalent concentrations. RNA was then reverse transcribed using the iSCRIPT cDNA synthesis kit (Bio-Rad) according to the manufacturer’s protocol. The resulting cDNA was diluted 1:5 in nuclease-free distilled H2O. RT-qPCR was performed in triplicate using the Power SYBR green PCR master mix (Thermo Fisher) and the Applied Biosystems QuantStudio 6 Flex RT-qPCR system. Primers used for RT-qPCR are listed in **Table S7**.

### RNA immunoprecipitation-qPCR and -seq

Native RNA immunoprecipitations were performed using the MAGNA RIP kit (Millipore) according to the manufacturer’s protocol. Briefly, protein A/G magnetic beads were incubated with 3 μg of antibody (METTL3: Abcam, ab195352; CSTF2: Bethyl Labs, A301-487A; CSTF2T: Bethyl Labs, A301-092A) or rabbit IgG antibody (Sigma-Aldrich, PP64B) per condition for 30 minutes at room temperature. ∼800 μg of lysate was incubated with magnetic beads at 4°C overnight with rotation. 10% input fractions were saved from the lysate sample (10% for RNA input, 10% for protein input). The magnetic beads were washed, and a 10% fraction was saved for immunoblotting verification, followed by proteinase K digestion, elution, and RNA extraction. Samples were analyzed by immunoblotting and RT-qPCR as described above. RNA enrichment was calculated as the percent of input in each condition. For sequencing, RNA-seq libraries were generated from both input and IP METTL3 RNA by preparing amplified cDNA using the Clontech Ultra Low Input RNA SMARTer mRNA amplification kit (Takara), followed by conversion of the double stranded cDNA into an Illumina sequencing library using the Kapa Hyper Prep Kit (Takara). The libraries were sequenced on the NovaSeq X Plus with PE read lengths of 2 x 150 base pairs. All samples yielded 67–83 million paired-end reads. Duplication rates ranged from 50–67%, consistent with expectations for amplified low-input RIP-seq libraries.

### METTL3 RIP-seq Analysis

All computational analyses were implemented as Snakemake workflows using GRCh38 as the reference genome with GENCODE v49 annotations^91^.

*Gene-level analysis*. Paired-end reads were aligned to the GRCh38 primary assembly using STAR v2.7.11b^92^ with GENCODE v49 splice junction annotations. Resulting alignments were filtered to retain reads with a mapping quality (MAPQ) ≥ 20 using samtools v1.22^93^. Gene-level read counts were quantified from filtered alignments using HTSeq v2.0.5^94^. Gene annotations were appended to count matrices using biomaRt querying Ensembl release 113^95^.

*Transcript-level analysis*. A decoy-aware Salmon v1.10.3^96^ index was built from GENCODE v49 transcript sequences with GRCh38 chromosomal sequences designated as decoys (k-mer size = 31; --gencode flag). Transcript-level abundances were estimated using selective alignment (--validateMappings) with GC bias (--gcBias) and sequence-specific bias (--seqBias) corrections; library type was set to IU. Per-sample outputs were imported into R v4.5.0 at isoform resolution using tximport (txOut = TRUE)^97^, and transcript annotations from Ensembl release 113 were merged with the resulting count matrices.

#### Differential binding analysis

Differential METTL3 binding was assessed using DESeq2^98^ (v1.48.2) in R at both gene and transcript resolution. Gene-level count matrices were generated from our STAR-aligned reads, while transcript-level counts were obtained from our Salmon quantification. Estimated counts were rounded to integers prior to analysis. We first identified genes that were significantly enriched by METTL3 in mock and ZIKV conditions. METTL3 binding was determined by comparing IP versus input samples using the Wald test. To ensure robust quantification, features were filtered to retain only those with a minimum of 10 reads in all samples within each condition. METTL3-bound genes and transcripts were defined as those significantly enriched in the IP fraction (IP/input Log_2_FC ≥ 0.58 and p_adj_ < 0.05).

Next, we analyzed differential METTL3 binding upon ZIKV infection. To identify infection-induced changes in METTL3 binding, differential enrichment between mock and ZIKV conditions was assessed using a ratio-of-ratios framework. Specifically, METTL3 binding was modeled as the change in IP vs input enrichment in ZIKV relative to mock (ZIKV_IP/Input_ – Mock_IP/input_). This was implemented in DESeq2 using a design formula of ∼ assay + condition + assay:condition, where the interaction term captures differential METTL3 enrichment between conditions. A likelihood ratio test (LRT) was performed by comparing the full model to a reduced model (∼ assay + condition) to identify features with significant changes in METTL3 binding. Features were filtered to retain those with a minimum of 10 reads across all samples. Genes and transcripts exhibiting increased METTL3 binding in ZIKV were defined as those with positive changes in IP vs input enrichment in ZIKV relative to mock (Log_2_FC ≥ 1 and p_adj_ < 0.05) and METTL3-bound in the ZIKV condition (IP/input Log_2_FC ≥ 0.58 and p_adj_ < 0.05). Conversely, decreased METTL3 binding was defined by negative changes in IP vs input enrichment in ZIKV relative to mock (Log_2_FC ≤ −1 and p_adj_ < 0.05) and METTL3-bound in the mock condition (IP/input Log_2_FC ≥ 0.58 and p_adj_ < 0.05). Features that were METTL3-bound in either condition but did not exhibit significant differences in IP vs input enrichment in ZIKV relative to mock were classified as unchanged.

### IsoformSwitchAnalyzeR

Isoform fractions were quantified using the IsoformSwitchAnalyzeR package^56^ (v2.8.0) in R. Transcript-level count matrices were generated from our Salmon quantification and imported as a transcript-by-sample matrix. A SwitchAnalyzeR list object was created using importRdata(), incorporating transcript counts and gene annotation from the GENCODE v49 reference. Annotation preprocessing options included removal of non-conventional chromosomes, collapsing transcript identifiers after periods, and correction of known StringTie annotation inconsistencies. Lowly expressed features were filtered using prefilter(), retaining genes and isoforms with expression ≥ 1 and restricting analysis to multi-isoform genes.

### GLORI-seq

GLORI treatment of purified mRNA was performed as described previously^36^. 20 ng of GLORI-treated mRNA was used for library preparation with the SMARTer Stranded Total RNA-seq kit (v3) Pico Input mammalian (Takara, 634487) according to the manufacturer’s protocol. The libraries were sequenced on the NovaSeq X Plus with PE read lengths of 2 x 100 base pairs.

### Data analysis for GLORI-seq

For calling high confidence m^6^A sites, we followed the GLORI-tools pipeline and filtering criteria described previously^36, 37^. First, files were trimmed and filtered as recommended, then paired-end reads were merged using PEAR^99^ before running GLORI-tools using the default parameters. To compare differential m^6^A methylation in ZIKV vs. mock, we took all high confidence sites detected in each sample and filtered them for sites present in at least 2 of the 3 replicates per condition in at least one condition (ZIKV or mock). For these filtered sites, we obtained A/G coverage information using the base-depth tool Perbase^100^ (Version 0.9.0). Obtained sites were filtered by depth>15 and sites were removed from further analysis if they did not meet the depth criteria in all the samples. Methylation ratio was recalculated for each site and differentially methylated sites (|Δm^6^A ratio|≥0.15, p<0.05) were detected with a binomial test using Methylsig as described previously^36, 101^.

### Direct RNA sequencing

Total cellular RNA was extracted from cells using TriZOL (Thermo Fisher) according to the manufacturer’s protocol and diluted to equivalent concentrations. polyA+ selection was performed using the NEBNext® Poly(A) mRNA Magnetic Isolation Module (NEB E3370) according to the manufacturer’s protocol. 250 ng of mRNA was prepared for nanopore sequencing using the Direct

RNA Sequencing Kit (Oxford Nanopore Technologies, SQK-RNA004) and sequenced on the PromethION24.

All samples were basecalled with Dorado (version 1.0.2) with flags --estimate-poly-a and --modified-bases-threshold 0.0. Modified bases were called with the following four modified base models: rna004_130bps_sup@v5.2.0_pseU_2OmeU@v1, rna004_130bps_sup@v5.2.0_m5C_2OmeC@v1, rna004_130bps_sup@v5.2.0_inosine_m6A_2OmeA@v1, and rna004_130bps_sup@v5.2.0_2OmeG@v1.

The basecalled reads were aligned to the UCSC hg38 reference genome (https://hgdownload.soe.ucsc.edu/goldenPath/hg38/bigZips/) with minimap2 (version 2.28) (10.1093/bioinformatics/bty191). Alignments were converted to sorted bam files with samtools (version 1.21) (10.1093/bioinformatics/btp352). Modification status was summarized by genomic position with modkit version 0.5.0 (https://github.com/nanoporetech/modkit). Gene-level aggregations were performed with custom scripts (https://github.com/Theo-Nelson/horner-zika). Transcript-level aggregations were performed with modulator v1.0.0.

### meRIP-qPCR

For each sample, 15 μg of total RNA was spiked with 0.1 fmol of positive control (m^6^A-modified *Gaussia* luciferase RNA) and negative control (unmodified *Cypridina* luciferase RNA) supplied with the EpiMark *N6*-methyladenosine Enrichment kit (NEB). meRIP was then performed as described previously^22^. Following meRIP, cDNA from the input and immunoprecipitated RNA fractions was generated and analyzed by RT-qPCR as described above. The relative m^6^A level for each transcript was calculated as the percentage of input under each condition, and the percent change of enrichment was calculated with siCTRL samples normalized to 100%.

## Notes

### Competing Interest Statement

CEM is a co-founder of Biotia.

### Summary of Updates

New figures: Figure 3, Figure 5, Figure 6, Figure 7, Figure 8; new supplementary files: Table S6, Table S7; title and abstract updated; manuscript updated to incorporate new results and discussion from Figures 3, 5-7; methods updated.

